# Reduction of cortical pulling at mitotic entry facilitates aster centration

**DOI:** 10.1101/2023.03.21.533625

**Authors:** Anne Rosfelter, Ghislain de Labbey, Janet Chenevert, Rémi Dumollard, Sebastien Schaub, Zoltan Machaty, Lydia Besnardeau, Céline Hebras, Hervé Turlier, David Burgess, Alex McDougall

**Affiliations:** Laboratoire de Biologie du Developpement de Villefranche-sur-mer, Institut de la Mer de Villefranche-sur-mer, Sorbonne Université, CNRS, 06230 Villefranche-sur-mer, France; Center for Interdisciplinary Research in Biology (CIRB), Collège de France, CNRS, INSERM, Université PSL, Paris, France; Department of Biology, Boston College, Chestnut Hill, Massachusetts, USA

**Keywords:** Sperm aster, Mitotic apparatus, Nuclear migration, Cell cycle

## Abstract

Although it has been studied for more than a century, the question of how one cell divides into two equal parts is still not fully resolved. Zygotes have provided much of the mechanistic insight into how the mitotic apparatus finds the center of the cell since the centrally-located mitotic apparatus is created from a large sperm aster that forms at the cortex and thus far from the zygote center. Here we show that in ascidians, the sperm aster extends throughout the cytoplasm during interphase yet remains located near the cortex and does not migrate towards the zygote center. It is only at mitotic entry, when the sperm aster has duplicated and the mitotic apparatus is being assembled, that most of the migration and centration occurs. This temporal pattern of centration behavior is mirrored by primate zygotes (including human). The current mechanisms of aster centration include cytoplasmic pulling that scale with microtubule (MT) length, MT pushing against the proximal cortex or MT-based cortical pulling. However, it is not yet known whether and how these 3 mechanisms are coordinated to prevent aster migration during interphase and trigger migration at mitotic entry. By monitoring quantitatively all three mechanisms (cytoplasmic pulling, pushing and cortical pulling) we have discovered that cortical pulling is switched off as the zygote enters mitosis while both cytoplasmic pulling and proximal cortical pushing remain active. Physical simulations could recapitulate both the static and migratory aspects of sperm aster and mitotic apparatus behavior. We therefore surmise that the reduction in cortical pulling at mitotic entry represents a switch that allows proximal cortical pushing forces and cytoplasmic pulling forces to center the nascent mitotic apparatus.

**Graphical abstract:** 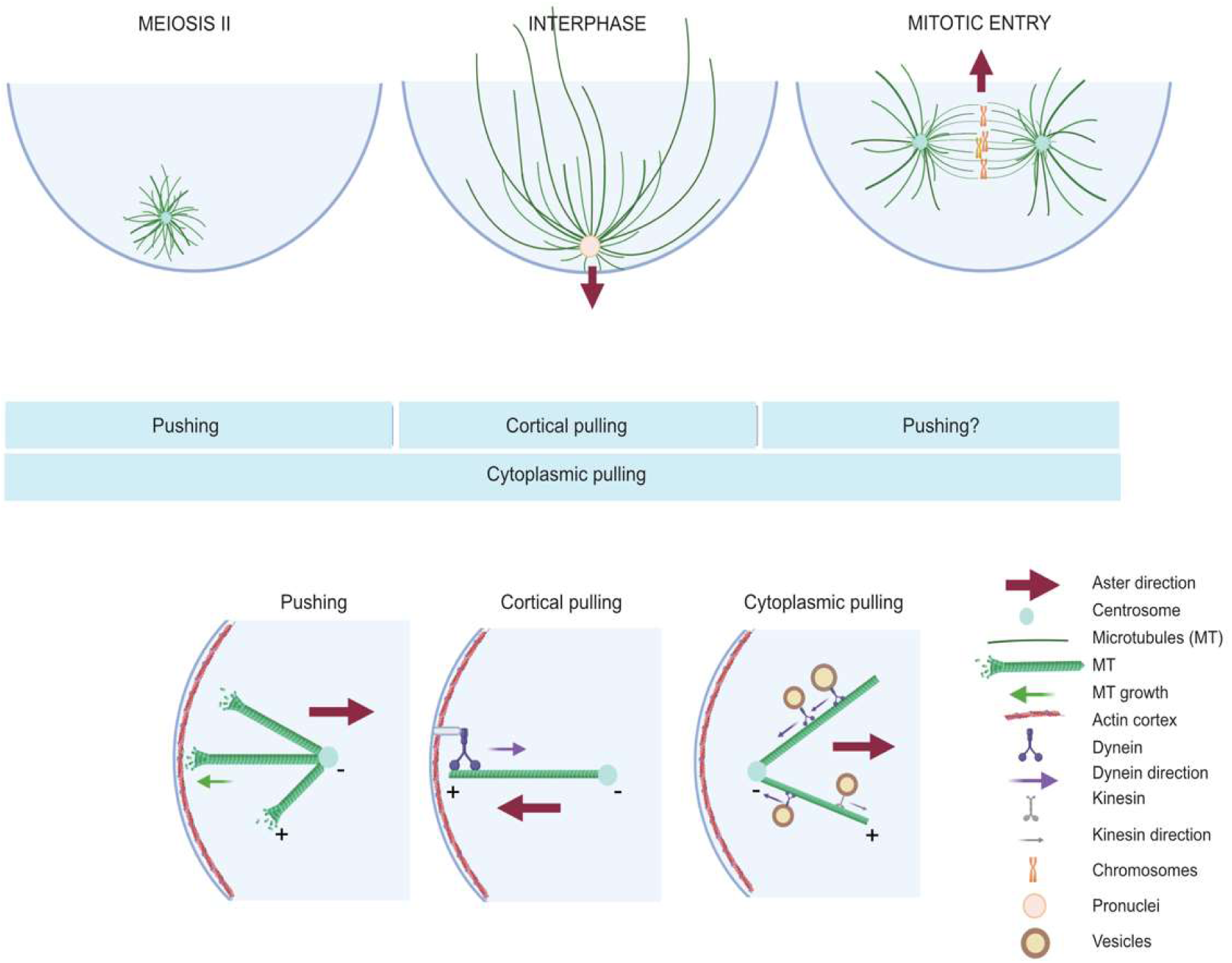

**Highlights:**

- Sperm aster/mitotic apparatus centration occurs at entry into first mitosis
- MT-based cortical pulling is active during interphase and switched off at mitotic entry
- Loss of cortical pulling at mitosis entry facilitates centration of the aster
- MT-based cytoplasmic pulling is active during both interphase and mitosis
- Agent-based simulations advocate the need for cytoplasmic pulling, a switch in cortical pulling and a minor role of pushing for aster centration at mitotic entry.

## Introduction

Asters are intracellular structures of microtubules (MTs) radially organized around a microtubule organizing center (MTOC), generally a centrosome (Bornens, 2012), or self-organized in presence of molecular motors (Mitchison and Field, 2021; Nédélec et al., 1997). The cell may contain a single aster, as in the case of the sperm aster, or two asters following centrosome duplication, which localize at each pole of the mitotic spindle (Meaders and Burgess, 2020). Asters are involved in many essential functions in the cell including intracellular trafficking and organization (Hamaguchi 1986, Smyth 2015), polarity establishment (Wodarz, 2002), guidance of nuclei (Reinsch and Gönczy, 1998), and determination of the cleavage axis during cell division (Devore et al., 1989; Strome, 1993). Although we have known for more than a century that the protoplasmic mass of the cell becomes equally segregated to the two daughter cells following cell division (Hertwig, 1884), the precise details of the mechanism of migration and centration of the mitotic apparatus is still not fully resolved.

Elegant experiments in amphibians, echinoderms, and *C.elegans* zygotes, as well as in yeast and *in vitro* studies, have reported three possible mechanisms of how MTs can generate forces to displace nuclei and whole asters/MTOCs. One mechanism is based on cytoplasmic pulling (Li and Jiang, 2018; Minc and Piel, 2012; Minc et al., 2011; Pelletier et al., 2020; Wühr et al., 2009), one on cortical pulling (Grill et al., 2001; Kotak and Gönczy, 2013; Redemann et al., 2010), and one on MT pushing (Garzon-Coral et al., 2016; Laan et al., 2012; Meaders and Burgess, 2020; Tran et al., 2001). During centration of the sperm aster cytoplasmic pulling exerts forces on the aster by taking advantage of the transport of organelles that travel on the MTs (Kimura and Kimura, 2011; Tanimoto et al., 2016). A cargo moving towards the MTs minus-end at the centrosome experiences an opposing drag force from the cytoplasm (Longoria and Shubeita, 2013; Palenzuela et al., 2020). Cytoplasmic pulling force thus scales with MT length as more cargoes cover longer MTs (De Simone et al., 2018; Kimura and Onami, 2005). Cortical pulling occurs when MTs contact a minus end directed molecular motor localized on the plasma membrane (Laan et al., 2012) often via interaction with cortical LGN/Pins/GPR1-2 and NuMA/Mud/Lin-5 complexes (Pietro et al., 2016). Such minus-end directed motor activity, supported by the rigidity of the actin cortex, can pull on MTs bringing the whole aster and mitotic apparatus (notably during asymmetric cell division) towards the cell membrane (Grill and Hyman, 2005; Grill et al., 2001; Kiyomitsu, 2019). Finally, due to the inherent dynamic instability of MTs (Burbank and Mitchison, 2006) and their polymerization against the membrane and its actomyosin rich cortex (Rosenblatt et al., 2004), MTs can exert a pushing force in the opposite direction to MT growth (Sulerud et al., 2020) and thus lead to displacement of asters away from the cell surface (Meaders and Burgess, 2020). Such force generation may be limited however by MT buckling when the MTs exceed about 20µm in length (Dogterom et al., 2005), although this would be more difficult to interpret if short branched MTs were also present (Field and Mitchison, 2018; Petry et al., 2013).

Striking examples of sperm aster or mitotic apparatus centration occur following fertilization and have been studied in a number of species. Following fertilization a sperm aster forms from the sperm-derived centriole (in many species) at the site of sperm-egg fusion and thus near the plasma membrane and as a consequence far from the center of the fertilized egg (Ishihara et al., 2014). Interestingly however, the cell cycle phase during which aster migration occurs towards the center of the fertilized egg varies depending on species and is independent of the timing of fertilization. For example, sperm aster centration occurs during first interphase in sea urchin immediately after fertilization (Minc et al., 2011) and in mouse oocytes during interphase following exit of meiosis II (Scheffler et al., 2021) whereas it occurs during first prophase in *C.elegans* (Gönczy et al., 1999). In *C. elegans* the sperm aster remains close to the cortex during interphase. In human oocytes (Asch et al., 1995) as well as those of other primates (Hewitson and Schatten, 2002; Simerly et al., 2019) centration also occurs at entry into mitosis (Asch et al., 1995; Simerly et al., 2019) when the sperm aster has duplicated to become the first mitotic apparatus. It is currently unknown how centration of these highly asymmetrically-positioned sperm asters in primates is 1) prevented during interphase and 2) triggered at mitotic entry.

To understand the mechanism of sperm aster/mitotic apparatus centration we analyzed the fertilized eggs of ascidians, which are a sister group to the vertebrates (Delsuc et al., 2006). As in mammals, fertilization occurs during meiotic metaphase in the ascidian (Meta I for ascidians, Meta II for primates/humans). We discovered that the sperm aster in ascidians behaves in a similar manner to those in primates/humans: the sperm aster remains cortical during interphase and migrates as a duplicated mitotic apparatus at mitotic entry. By combining experiments and numerical simulations, we investigated how the asymmetrically positioned sperm aster is prevented from migrating towards the center of the zygote during interphase and in addition how centration is triggered at mitotic entry. We found that cortical pulling is switched off at mitotic entry facilitating aster centration at mitotic entry when both cytoplasmic pulling and pushing are active.

## Results

### Aster migration in *Phallusia mammillata* correlates with the cell cycle

To monitor sperm aster and then mitotic apparatus migration while observing cell cycle stages during the first cell cycle, oocytes of *Phallusia* were co-injected with mRNAs encoding MT-binding protein Ensconsin (Ens::3GFP) and Tomato-tagged histone H2B (H2B::Tom). These oocytes were fertilized and imaged from 10 minutes post fertilization (mpf) until cytokinesis. We observed that the aster changes shape, size and position throughout the cell cycle as detailed in the next paragraph (Figure 1A, Movie 1). These changes were associated with cell cycle stages based on recognizable events: polar body (PB) extrusions, pronuclei formation, female pronucleus (PN) migration, nuclear envelope breakdown (NEB), metaphase and anaphase (Figure 1C, 1D).

**Figure 1.**
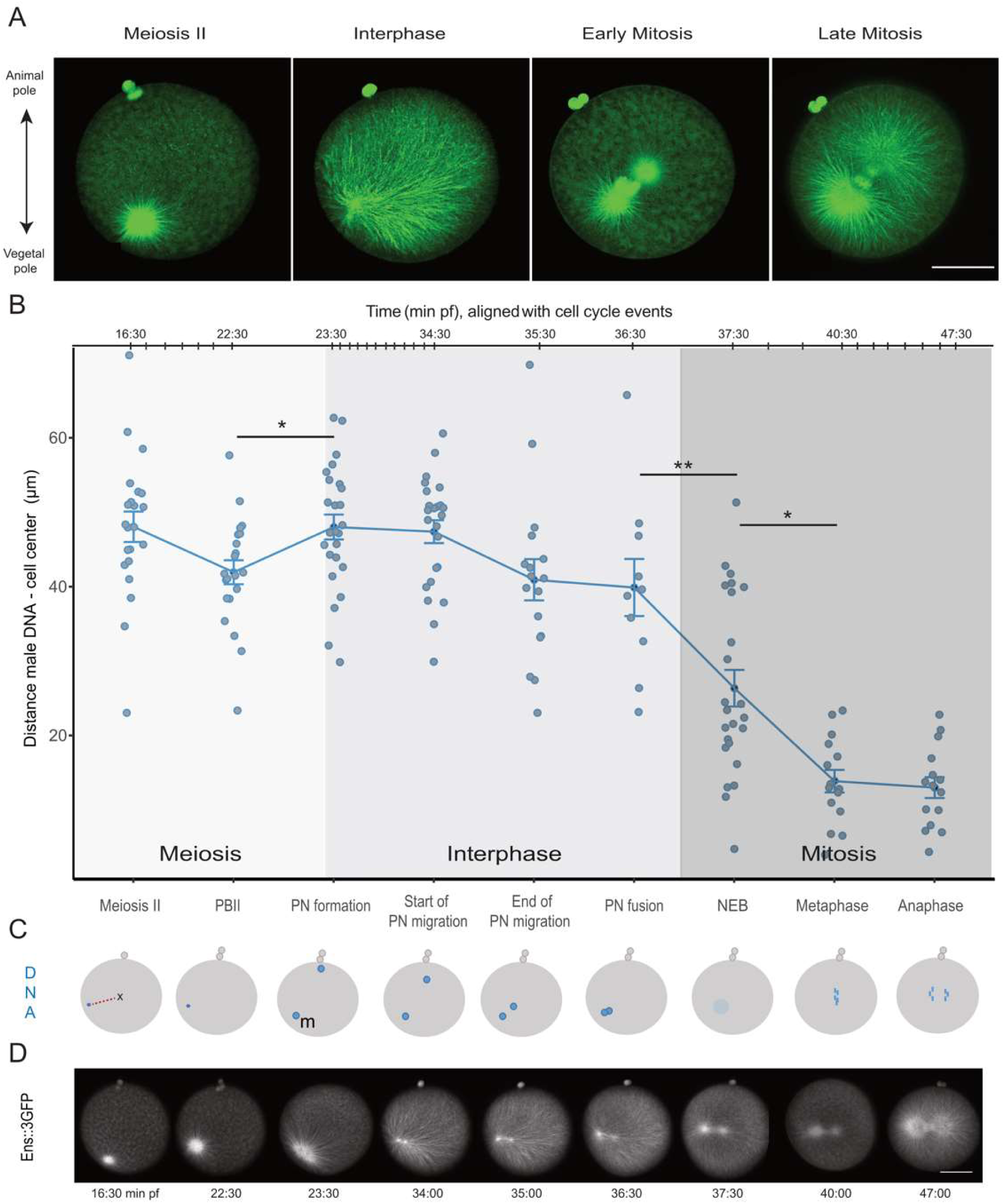
Migration of the sperm aster correlates with the cell cycle. A.Confocal z-sections from a xyzt series showing microtubules in a live zygotes labeled with Ens::3GFP. Meiosis II, interphase and mitosis (early and late) are displayed. See Movie 1. B. Quantification of the distance (in µm) between the male DNA and cell center measured in 3D at each event (see Materials and Methods) during the first cell cycle in 28 live zygotes. Graph showing the distance of the male DNA from the cell center (in µm) at each cell cycle event. The x-axis shows the cell cycle phases corresponding to the time points measured (bottom of the graph), and the timing of the zygote shown in Fig 1D relative to cell cycle event is indicated (top of the graph). The time scale is not linear but adjusted to spread evenly all cell cycle events. The graph is colored according to the cell cycle phases: meiosis II (light grey), interphase (medium grey) and mitosis (darker gray). Error bars represent SEM. Paired t-test adjusted with bonferroni, p-value ≤ 0.05 (*), ≤ 0.01 (**), ≤ 0.001 (***), p-value > 0.05 (ns). C. Schematic representation of the zygotic events that mark the first cell cycle and how they associate with DNA displacement in the cell. X in the first drawing marks the cell center from which distances are measured. Male pronucleus (m). D. Confocal images from a timelapse movie of a zygote with microtubules labelled with ENS::3GFP corresponding to the schematic in C. See also Movie 1. Scale bars in A and D are 50µm.

In ascidians, during the completion of meiosis, the sperm aster grows in the vegetal hemisphere, on the opposite side of the fertilized egg from the meiotic spindle which defines the animal pole (Roegiers et al., 1995). During meiosis I, the aster is in the egg cortex (Dumollard and Sardet, 2001) whereas at meiosis II (7 to 20 min after fertilization), the aster is spherical and located a few microns (5-10 µm) from the egg cortex (Fig 1A « meiosis II »). Upon entering interphase, when the pronuclei form (Figure 1A “interphase”), the aster remains close to the plasma membrane in the vegetal hemisphere and MTs elongate. Later in interphase, the MTs reach the opposite side of the zygote thus allowing the capture of the female PN formed at the animal pole (Movie 2). This leads to a highly asymmetric aster with long MTs toward the cell center and animal pole, and shorter MTs towards the vegetal pole. The centrosome then duplicates and the two resulting centrosomes position on each side of the male PN which gives the aster an oblong shape extended along the vegetal cell membrane (Figure 1A, “interphase”). Finally, in prophase, which as in subsequent embryonic mitoses occurs 3-4 minutes before NEB (Dumollard et al., 2013), two asters with short MTs are linked by a central mitotic spindle (Figure 1A “early mitosis”). On the time series shown in Fig. 1A the aster does not move between meiosis and interphase whereas between interphase and early mitosis the aster migrates towards the cell center. The separation of the two centrosomes and the formation of the nascent mitotic apparatus both occur during the studied time period; for clarity purposes we refer to the globality of the aster and subsequent spindle migration as “aster migration”.

Aster migration was quantified by measuring the distance between the cell center and the male DNA from 28 different embryos at several cell cycle stages (Figure 1B, Figure S1A). In the embryos pooled in the dataset the DNA was labelled either by expressing H2B::Tomato or H2B::Venus or by staining with Hoechst in live zygotes. The first noticeable event in the course of aster migration was the movement of the aster toward the cell cortex upon entry in interphase. Indeed, between the extrusion of the second polar body and the formation of the pronuclei, the aster moved away from the cell center (Fig. 1B). Then, during interphase (9 minutes duration), the distance between the male DNA and the cell center did not change significantly (Fig. 1B) suggesting that the aster did not significantly migrate. The main movement of migration occurred after pronuclei (PN) fusion, at mitosis onset. The distance of the DNA from the cell center decreased between the PN fusion and NEB and further declined between NEB and metaphase (Fig. 1B). Thus, after its initial growth in meiosis which displaces the aster slightly from the cortex, the sperm aster follows three phases of migration: a first movement brings the aster back to the cortex during which the aster changes shape (between 22 min. and 23 min. in Fig. 1B and 1D), then the aster does not move relative to the cell center during interphase (between 23min. and 35 min. in Fig 1B, 1D), and finally the aster starts centering at mitosis entry (after 35 min. in Figure 1C, 1D). This temporal sequence of aster migration was confirmed by the quantification of fixed samples from batches of zygotes sampled every 10 min and immuno-stained to label MTs and DNA (Figure S1B).

### Aster migration requires mitosis entry

The main movement of migration starts at mitosis entry (Figure 1). Therefore, we investigated the causal role of mitosis entry on the aster migration. We monitored DNA position as a read out for aster migration and perturbed the cell cycle by injecting oocytes with p21 protein, a cyclin-dependent kinase (CDK) inhibitor (Figure 2). The presence of p21 perturbs CDK activity (Levasseur et al., 2007) and thus prolongs interphase by delaying entry into first mitosis. The injection was considered successful when interphase (the time between PN formation and NEB) lasted more than twice the mean duration in control zygotes or greater than 24 min (Table S1).

**Figure 2.**
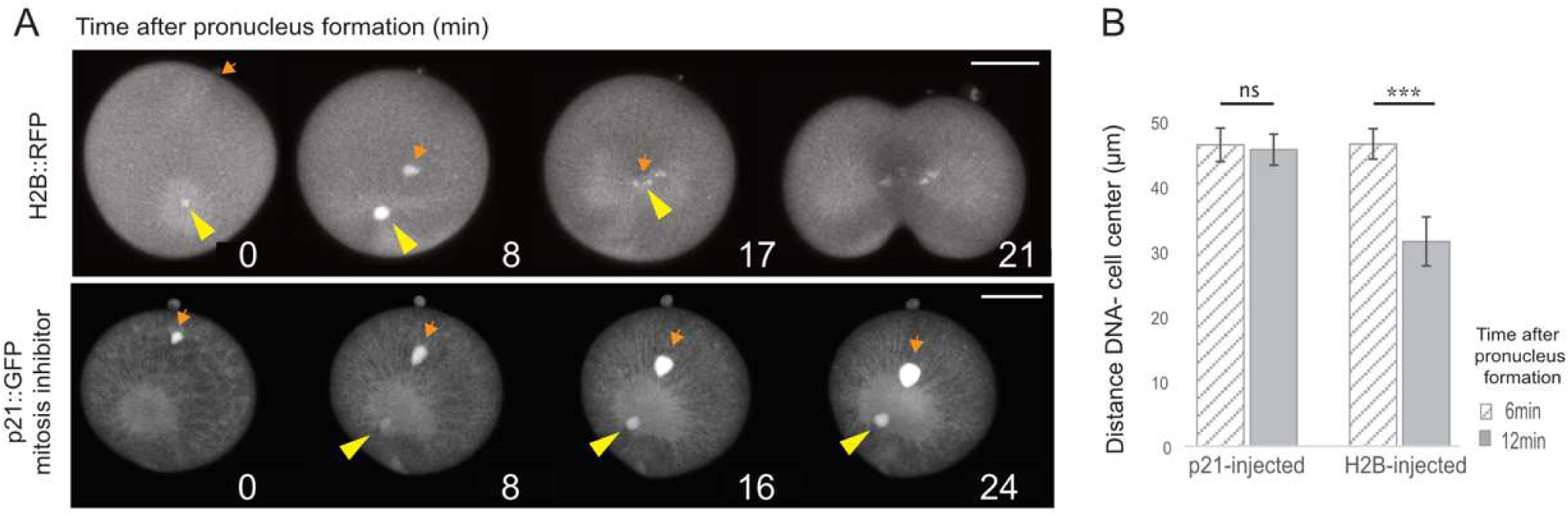
Entry into mitosis triggers aster migration. A. Representative time-lapse series of zygotes expressing either H2B::RFP (top panel) or injected with the cyclin-dependent kinase inhibitor p21::GFP protein (bottom panel). The male PN position, reflecting the aster position is shown by the DNA signal in H2B::RFP zygote and by the nuclear localization of p21::GFP in p21 injected zygotes. Male PN is indicated with a yellow arrowhead, female PN with an orange arrow. Time in minutes is indicated on each panel. Scale bars are 50µm. B. Quantification of the distance between the male DNA and the cell center (in µm) during interphase (hatched bars, 6 min after PN formation) and during mitosis entry (shaded bar, on average at 12 min after PN formation in controls) in p21-injected zygotes and in zygotes with an unaltered cell cycle. Error bars represent SEM. Paired t test. n=17 p21-injected embryos and n=17 control embryos.

In zygotes with an unaltered cell cycle (Figure 2A, top panel) the pronuclei formed, the female PN (orange arrow) migrated and reached the male PN (yellow arrowhead), then the mitotic apparatus migrated towards the cell center during mitosis (t=17 min post PN formation). In the p21-injected zygotes with prolonged interphase (Figure 2A, bottom panel), the male and female pronuclei formed and the female PN migrated toward the male PN until it reached the centrosome (Figure 2, t=16 min) or the male PN. However, the aster and the male PN never migrated, the aster remained asymmetric and uncentered and astral MTs continued to grow (Figure 2, t=24 min).

Quantification of the aster migration in the two conditions revealed that in control embryos the distance between the cell center and the male PN decreased from 45+/- SEM µm to 30 +/- SEM µm from interphase to mitotic entry (Fig 2B). In contrast, during the same timing of interphase in p21-injected zygotes the distance remained at 45 +/-SEM µm, showing an absence of aster migration. Thus, prolonging interphase prevents DNA centration and aster migration.

### Cytoplasmic pulling is constant during the cell cycle

Given that the aster specifically centers at mitotic entry, we wondered what could cause the absence of migration in interphase. The three mechanisms able to move the aster are cytoplasmic pulling (Li and Jiang, 2018; Minc et al., 2011; Wühr et al., 2009), cortical pulling (Grill et al., 2001; Kotak and Gönczy, 2013; Redemann et al., 2010) and cortical pushing (Garzon-Coral et al., 2016; Laan et al., 2012; Meaders and Burgess, 2020). Since the aster is highly asymmetric and has long MTs directed towards the animal hemisphere throughout the interior of the zygote during interphase (Figure 3A, Fig. 1A), one could expect to observe an aster migration by cytoplasmic pulling (Minc et al., 2011). Thus, we first tested whether the absence of migration during interphase was caused by a lack of cytoplasmic pulling. We examined whether these long interphasic MTs accumulated or transported organelles towards the aster center, which would indicate minus-end directed transport and cytoplasmic pulling (Kimura and Kimura, 2011). The accumulation of organelles was tested by characterizing minus-end directed transport in different contexts. First, during interphase the sperm aster captured the female PN which migrated towards the sperm aster center and male PN (Figure 3B). When imaging endoplasmic reticulum during interphase, most of the ER was accumulated around the aster (Fig 3C). Finally, and in order to characterize more precisely the dynamics of organelle movement towards the sperm aster, we imaged the transport of vesicles endocytosed from the plasma membrane (labeled with the membrane dye Cell Mask Orange) and measured their movement towards the sperm aster in the zygote (Movie 3A and 3B, Figure3D and Figure S2). The presence of minus-end directed transport of the female PN, the ER, and the endocytosed vesicles along the MTs during interphase demonstrated the presence of cytoplasmic pulling forces during interphase. We then wondered if cytoplasmic pulling may be weaker in interphase than in mitosis perhaps explaining the lack of sperm aster movement during interphase versus mitosis. As a proxy for cytoplasmic pulling, we assessed whether the vesicle traffic changed throughout cell cycle. We imaged endocytic vesicles at high-speed (1 image/sec) in one confocal plane and made time projections to visualize vesicle trajectories over the course of 3 minutes (Fig 3E). We categorized the vesicles as vesicles moving towards the aster, moving away from the aster or static (Figure 3E, F, Figure S2). The projection of vesicle tracks in an interphasic versus mitotic zygote illustrated that there is no major difference in interphase versus mitosis (Figure 3E). We then compared the speed and total transport of the endocytosed vesicles (Figure 3F). The speed measured is the mean speed of the vesicles on their total trajectories, and the total transport represents the sum of the distance traveled by all vesicles towards the aster center during the 3 min period. No statistically significant difference between interphase or mitosis was found for vesicle speed or the distance travelled (Figure 3F and Figure S2).

**Figure 3.**
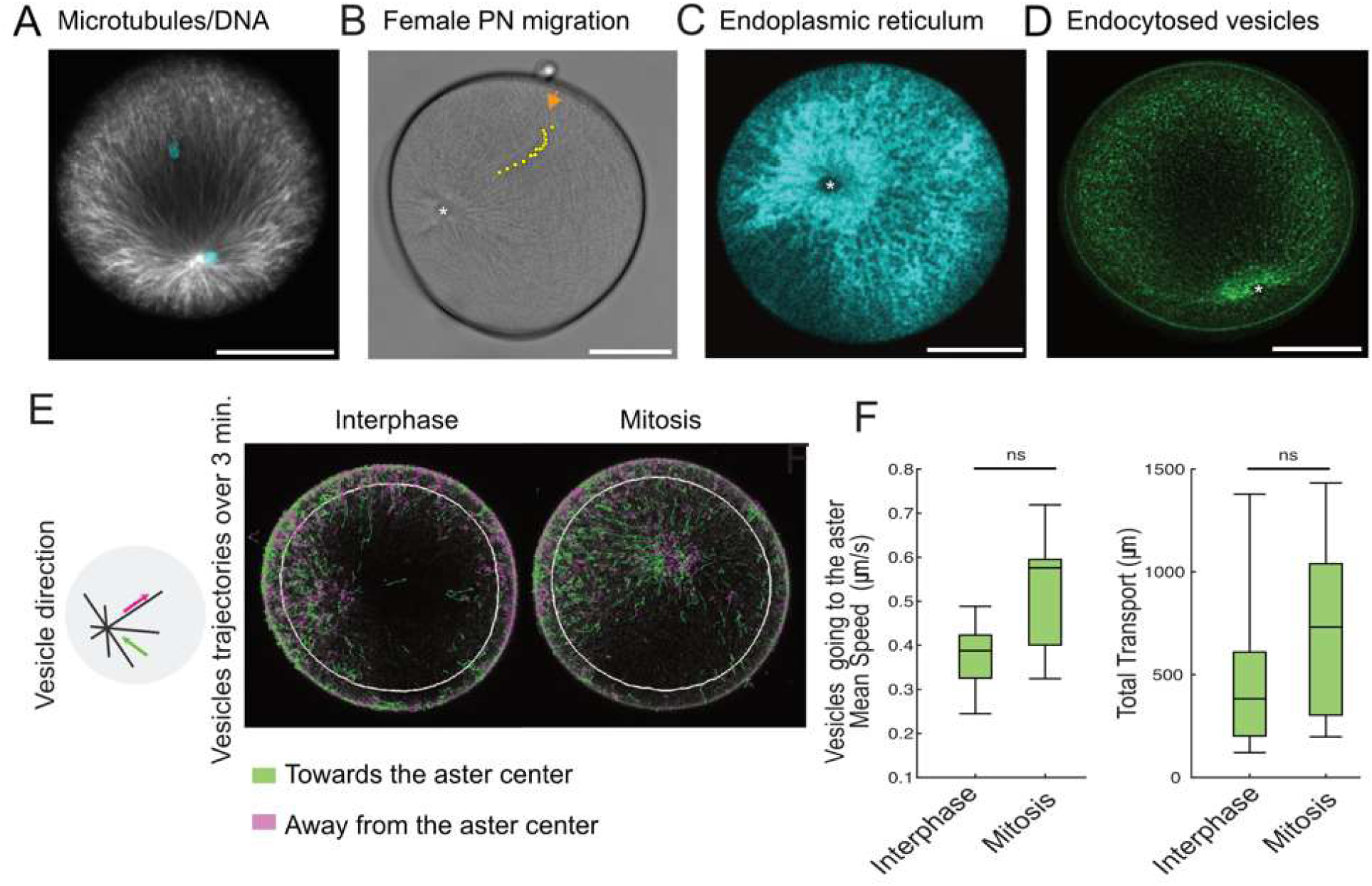
Characterization of minus-end directed transport in zygotes. A. Confocal image of a zygote fixed in interphase and immuno-labeled with the anti-tubulin antibody DM1a (white), and stained for DNA with Hoechst (blue). Long microtubules crossing the whole zygote are observed during interphase. B. Time projection of a bright field movie of a zygote showing migration of the female PN towards the male PN during interphase. Yellow dots show the female PN position at several time intervals. The male pronucleus is indicated by a white asterisk (*). An orange arrow indicates initial position of the female PN. See also Movie 2. C. Confocal image showing ER distribution in a zygote expressing Venus::Reticulon (cyan). Note that most of the ER is accumulated around the aster at this stage. The male PN is indicated by a white asterisk (*). D. Confocal image of a zygote stained with Cell Mask Orange showing endocytic vesicles accumulated at the sperm aster (green). The male PN is indicated by a white asterisk (*). Also see Movies 3A and 3B. All scale bars are 50µm. E. Left, drawing explaining vesicle trajectories (left panel). Vesicles were classified as moving towards the aster center (retrograde in green), or away from the aster center (anterograde in magenta). Right, time projections over 3 min of a zygote treated with Cell Mask Orange (CMo) in interphase and in mitosis showing the tracks of vesicles endocytosed from plasma membrane. Tracks of anterograde vesicles are in magenta, retrograde vesicles in green. The white circle denotes the area in which vesicle were quantified. F. Quantification of the speed (in µm/sec) and total transport of CMo vesicles going towards the aster. Vesicles were imaged during 3 minutes of interphase and during 3 minutes of mitosis. Speed is measured over the complete vesicle trajectory, including pauses (left graph) and Total transport is the cumulated distance travelled by all retrograde vesicles in direction of the aster center (right graph). Box plot central mark indicates the median, bottom and top edges of the box indicate the 25th and 75th percentiles. The whiskers extend to the most extreme data points without considering outliers. Paired Wilcoxon test. n=8 embryos.

To conclude, we have demonstrated that there is retrograde transport and hence, active cytoplasmic pulling forces during interphase, and that the organelle transport is not significantly different in interphase and mitosis. This indicates that another mechanism must account for the difference of aster migration behavior between interphase and mitosis, and especially, for the lack of aster migration in interphase.

### Cortical pulling is strong in interphase and weak in mitosis

The asymmetric configuration of the sperm aster in interphase together with the accumulation of organelles at the aster center (Figure 1A, “interphase”) should favor an interphasic aster migration by cytoplasmic pulling in the direction of the longer MTs (Kimura and Kimura, 2011; Tanimoto et al., 2016). Since there was no aster migration despite the presence of cytoplasmic pulling, we hypothesized that some mechanism must prevent aster migration. A candidate mechanism to counteract the centering forces of cytoplasmic pulling is cortical pulling, which is able to off-center an aster (Laan et al., 2012). Cortical pulling occurs when minus-end directed motors bound to the cortex interact with MTs (Laan et al., 2012). The transport of the plasma membrane to the centrosome is prevented by the thick actin cortex which provides a support that resists membrane deformation. Hence, with an intact cortex, instead of bringing the plasma membrane towards the MTs minus-end, the molecular motors pull the centrosome to the plasma membrane (Figure 4A). We analyzed whether such cortical pulling forces would thus restrain the aster from migrating to the cell center.

**Figure 4.**
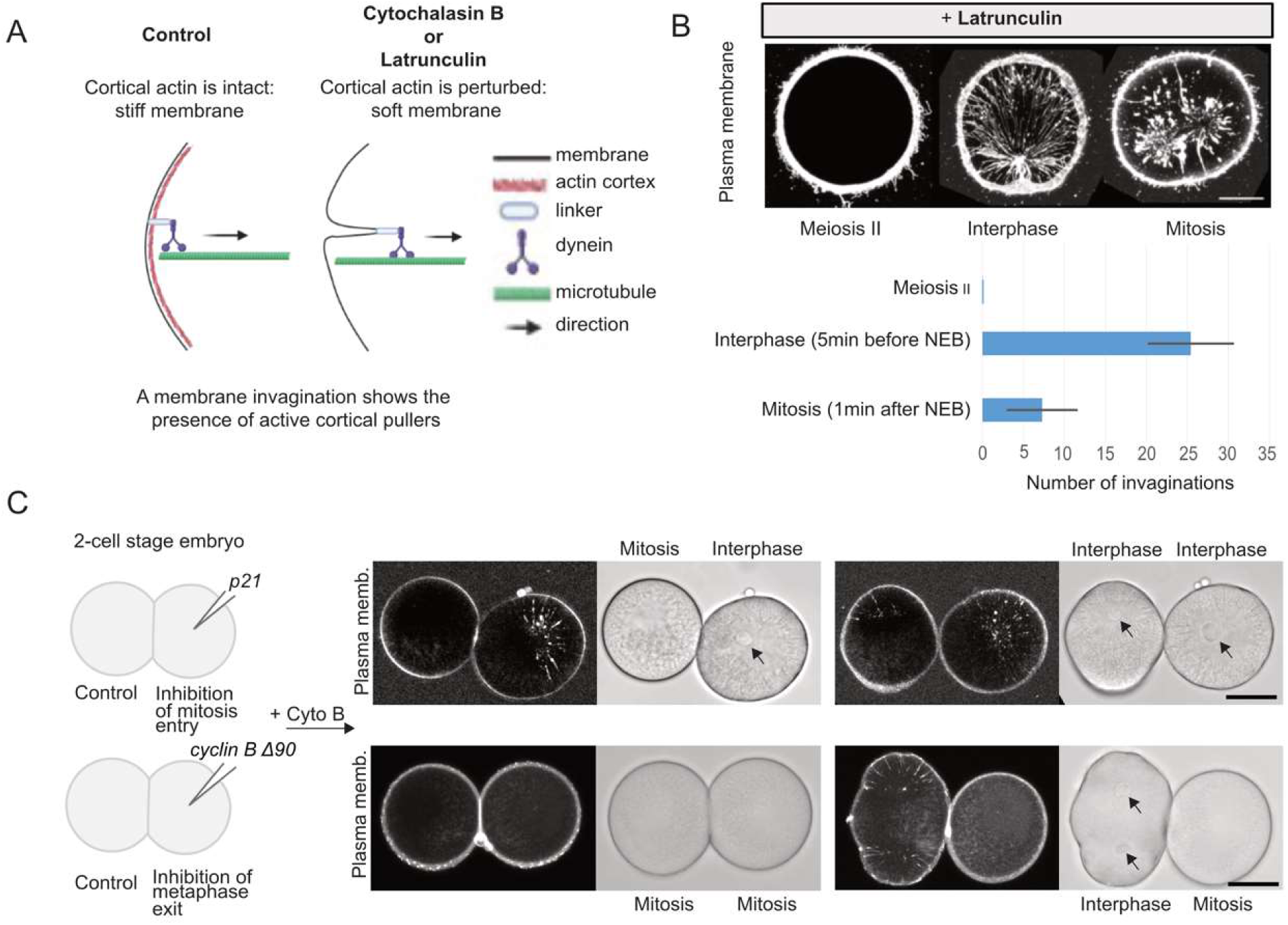
Cortical pulling is stronger in interphase. A. Schematic representation of the invagination assay that tests the presence of cortical pullers attached to the membrane. B. Top panel: Confocal images of the membrane invaginations in meiosis, interphase and mitosis, in a representative zygote treated with latrunculin and labelled with CMo. Images are a z-projection of 10 images from a 25 µm-thick z-stack of confocal images. Bottom panel: Number of invaginations in meiosis (meiosis II), in interphase (5 min before NEB) and in mitosis (1 min after NEB). Error bars represent +/- SD. n=7 embryos. Also see Movie 4. C. Schematic representation of the injection experiments to perturb cell cycle in one cell of 2 cell stage embryo. Plasma membrane was labelled by microinjection of PH::GFP mRNA into eggs prior to fertilization, then at the 2-cell stage 1 cell was injected with the indicated proteins. Images on left show plasma membrane label (white) in both the injected cell and the control sister cell. Images are a projection of 10 images from a 10µm-thick z-stacks of confocal images at different time points extracted from xyzt movies. The cell cycle stage of each cell is indicated above and below the BF images. Plasma membrane invaginations are observed only in cells in interphase (whether from control side or protein-injected side of the embryos). Black arrow indicates the presence of a nucleus. n=11 embryos for p21 and n=9 embryos for Δ90. Also see Movies 5A and 5B.

We examined cortical pulling over the zygote cell cycle using a membrane invagination assay (Godard et al., 2021; Redemann et al., 2010; Rodriguez-Garcia et al., 2018). We used two pharmaceutical compounds to disrupt the actin cortex: cytochalasin B or latrunculin B. Both perturbed the cortical resistance necessary to prevent membrane deformation. Thus, by labeling the membrane, invaginations of the plasma membrane towards the centrosome could be observed at sites of cortical pulling (Figure 4B, Figure 4C). Depolymerization of MTs with nocodazole completely inhibits such invaginations (Godard et al., 2021). In zygotes treated with latrunculin, the plasma membrane invaginations appear in a cyclic manner (Figure 4B, Movie 4). We determined the mean number of invaginations at different phases of the cell cycle (Figure 4B bottom). Invaginations were never observed at meiosis II (6 minutes before PN formation) while an average of 25 +/- 5 (SEM) invaginations was counted during interphase (5 minutes before NEB) that decreased to 7 +/- 5 (SEM) invagination at mitosis (1 minute after NEB) (Figure 4A, 4B bottom). These observations demonstrate that cortical pulling is correlated with the cell cycle as it is not active in meiosis and mitosis but very active in interphase.

To test the causality between the cell cycle phases and the presence of active cortical pullers, we performed the membrane invagination assay on 2-cell stage embryos in which one cell was either blocked in mitosis or stalled in interphase while the sister cell kept cycling and served as a non-injected control cell. To manipulate the cell cycle we injected either p21 protein, to delay entry into mitosis, or a truncated non-destructible form of cyclin B protein (Δ90-cycB) to prevent exit from metaphase (Figure 4C and Figure S3) (Levasseur and McDougall, 2000). As for the zygotes (Figures 4B) the non-injected cell displayed invaginations in interphase while none were visible in mitosis. We found that in p21-injected cells membrane invaginations were detected all throughout the prolonged interphase (Figure 4C top row and Movie 5A). On the contrary, in cyclin B Δ90 protein injected cells, no invaginations were visible during the prolonged metaphase, while the non-injected sister cell displayed invaginations during interphase (Figure 4C bottom row and Movie 5B). The absence or presence of plasma membrane invaginations as a function of time and cell cycle phase is displayed for each 2-cell stage embryo (Figure S3). In non-injected cells the invaginations disappeared around the time of NEB (stars in Figure S3), and reappeared about 10-15 min after NEB at mitosis exit. We conclude that cortical pulling is cell cycle dependent, it is up-regulated during interphase and down-regulated at mitotic entry, prometaphase and metaphase.

### Evaluation of the necessity of both cortical pulling and MT pushing against the cortex for aster migration

Cortical pulling is active in interphase therefore it could be responsible for preventing aster migration before mitotic entry. If cortical pulling is inhibited an early aster migration is expected. The inhibition of cortical pulling was performed by adding latrunculin which perturbs the actin cortex. This drug should also inhibit the pushing mechanism since it also requires an intact cortex. By examining aster migration in the presence of latrunculin, we evaluated the necessity of both cortical pulling and MT pushing against the cortex for aster migration. We found that when latrunculin was added during meiosis, the sperm aster was on average 60 µm away from the cell center and remained so for more than 10 minutes. Then the aster migrated toward the cell center during mitosis (Figure 5A, orange). In contrast, the aster in control DMSO-treated zygotes (blue in Figure 5A) was on average 40 µm away from the center and first migrated slightly towards the cortex at interphase entry (to 45 µm away from the center) and remained in position until mitotic entry, when the aster moved toward the cell center. The initial cortex-directed movement (just before PN formation) was not observed in latrunculin-treated zygotes (Figure 5A). Nevertheless, in both latrunculin and DMSO conditions, the aster migrated approximately 15µm (total distance) between PN formation and NEB (Figure 5B), suggesting that cytoplasmic pulling is sufficient to move the aster towards the cell center. Since latrunculin inhibited cortical pushing we suggest that the closer proximity of the sperm aster to the plasma membrane is caused by the lack of efficient cortical pushing in meiosis. At two minutes before NEB we noted a slower (though not significant) pace of migration in latrunculin-treated zygotes (Fig 5B) which suggests that an additional mechanism other than cytoplasmic pulling may support aster migration at mitotic entry in control condition.

**Figure 5.**
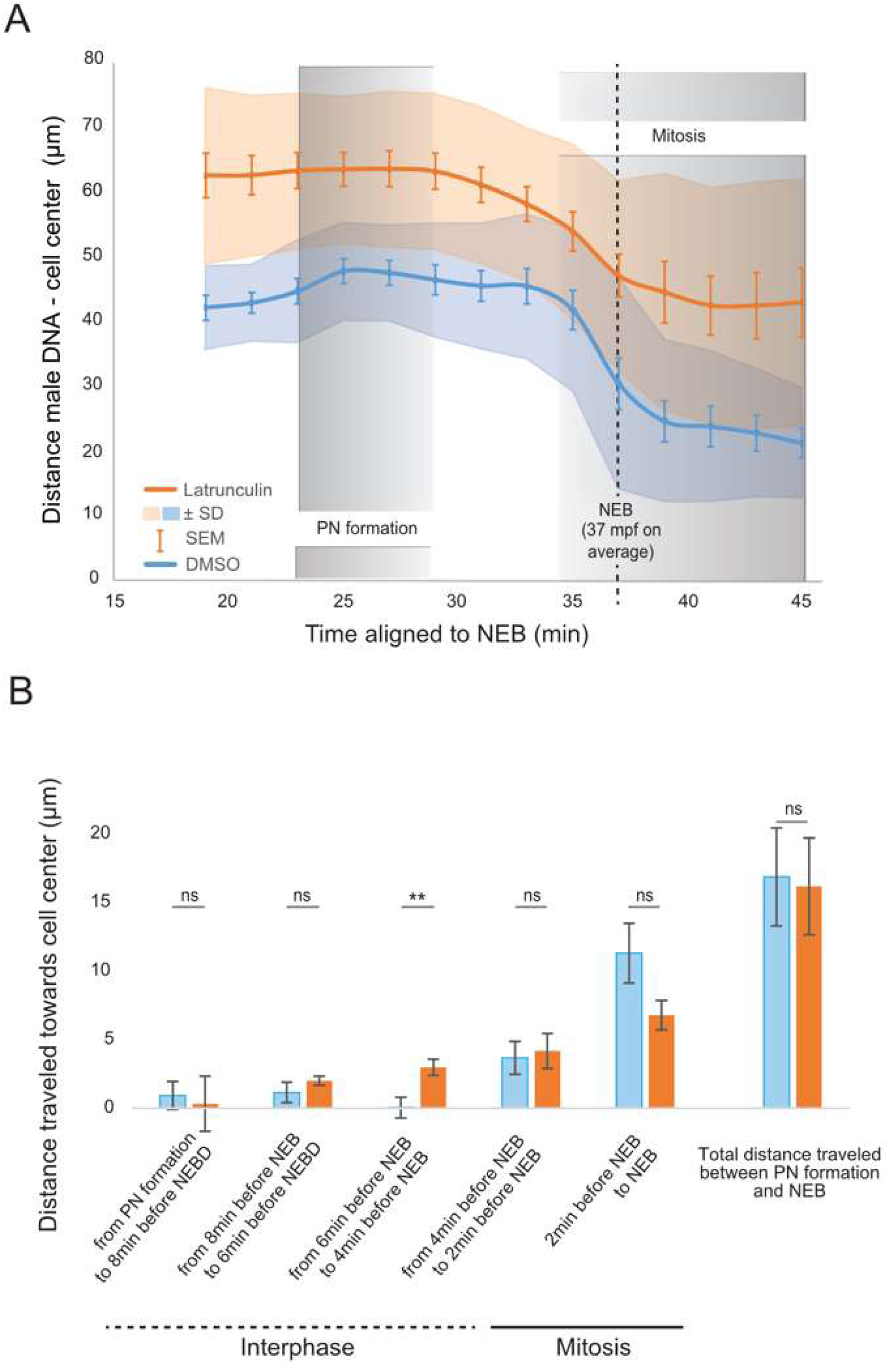
In absence of cortical pulling and pushing the aster migration is not advanced in time. A. Average distance (in µm) from DNA position to cell center over time. The distances are measured in two conditions; in presence of DMSO (control, blue curve, n=16) and in presence of latrunculin (orange, n=20), an actin polymerization inhibitor. To compute the average curve, individual curves were aligned with respect to the time of NEB, here represented by a dotted line. The time from PN formation to NEB is not equivalent in each embryo, thus PN formation is indicated as a span in gray shading. Mitosis is also indicated with a gray shading. The orange and blue shades represent +/- SD, Error bars = +/- SEM. B. Distance travelled by the sperm aster towards the cell center following latrunculin (orange bars) or DMSO (blue bars) treatment. All comparisons are not statistically different except for timepoint 6 to 4 min before NEB which corresponds to the time when the sperm aster is flattened against the cortex in interphase. Error bars = +/- SEM. Wilcoxon test.

To sum up, the absence of a clear early aster migration when cortical pulling is inhibited (Figure 5A, 5B) shows that, in addition to cytoplasmic pulling which can center the aster (Figure 5A), a second mechanism is involved at mitotic entry to permit the fast aster migration. Since the inhibition of cortical pulling is accompanied by the inhibition of pushing, we hypothesized that the pushing mechanism could be an actor of aster centration at mitotic entry, thus we sought to determine when the pushing mechanism is active.

### Cortical pushing is active in meiosis and mitosis

Following latrunculin treatment the aster was closer to the membrane than it was in control zygotes. We suggest that the closer proximity of the sperm aster to the plasma membrane caused by latrunculin could be explained by the lack of cortical pushing during meiosis II (Figure 5A). Indeed, we observed during meiosis II that the presence of an aster close to plasma membrane with a perturbed actin cortex creates thin membrane protrusions (i.e., a membrane deformation towards the exterior, Figure 6A). These deformations were not stable as they grew and shrunk (Movie 6) and co-localized with MTs (Figure 6B). This co-localization of MTs and plasma membrane as well as the dynamic nature of the protrusions suggested that the protrusions were caused by MT polymerization. It also supported the notion that an actin cortex was required for the pushing mechanism to move the nascent sperm aster away from the cortex. The protrusions were visible in meiosis, and in cases where the aster did not migrate during the cell cycle, thus permitting the observation of protrusions throughout the complete cell cycle. Protrusions were absent in interphase and present again just after NEB (Figure 6B, Movie 6). These deformations thus appeared to follow the cell cycle as do microtubules themselves, being more dynamic during mitosis but long and stable during interphase (Figure 1). These latter observations must however be taken with caution since the absence of protrusion in interphase could also be due to the presence of the nucleus that may prevent direct contact of MTs on the membrane.

**Figure 6.**
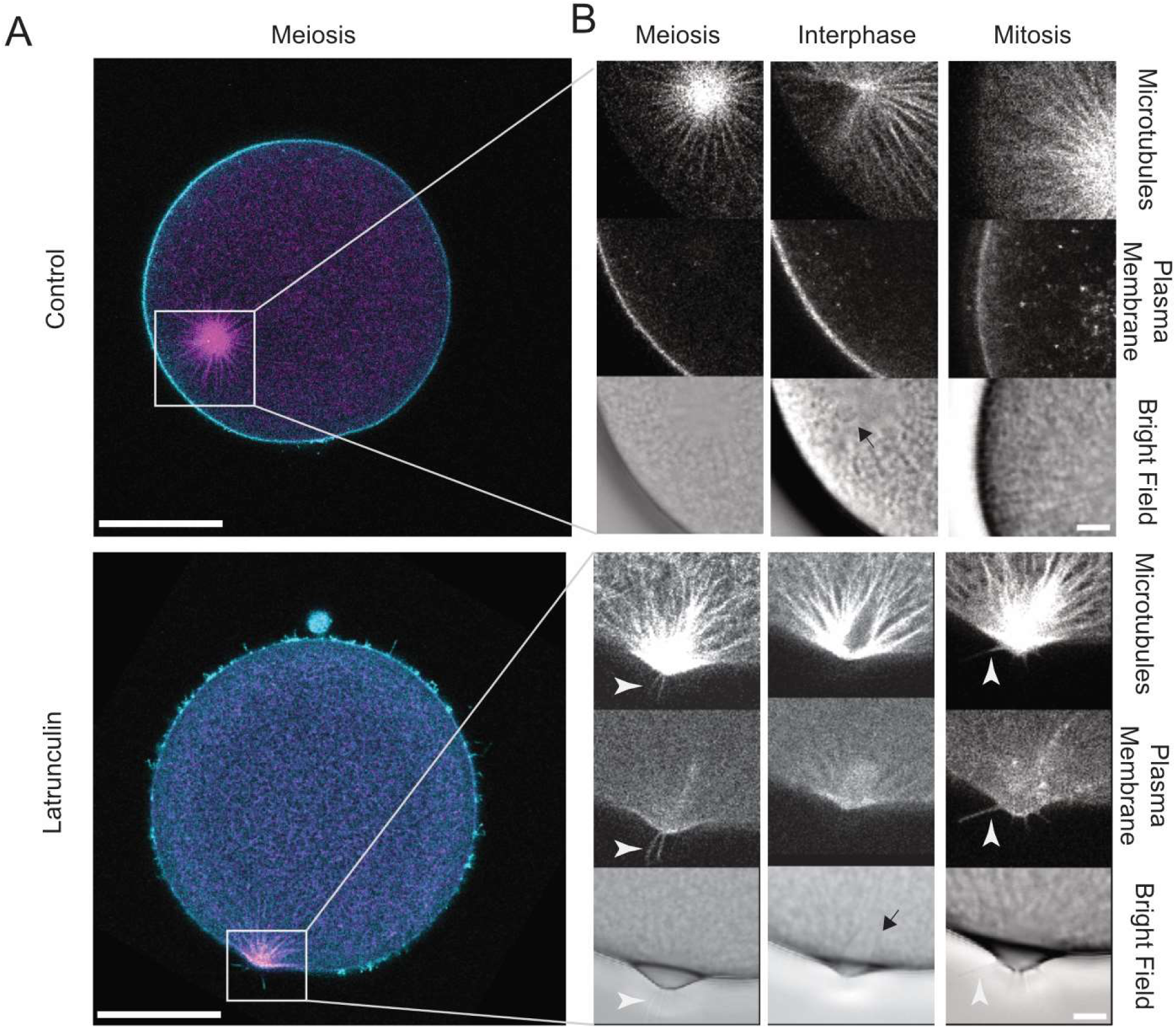
MTs push on the plasma membrane. A. Representative confocal images of a control zygote (top panel) and of a zygote treated with latrunculin (bottom panel), both in meiosis and expressing the microtubule marker Ens::3GFP and labeled with the plasma membrane dye Cell Mask Orange. Scale bar is 50µm. B. Close-up of A on the area containing the sperm aster and the closest piece of plasma membrane in meiosis (first column), in interphase (second column), and mitosis (third column) extracted from time-series of the same zygotes. Black arrows indicate the presence of a nucleus, white arrowheads indicate the presence of thin protrusions. Also see Movie 6. n=3 embryos out of the 9 where protrusions were seen in meiosis. Scale bar is 10 µm.

### A computer simulation to challenge the contribution of cortical pulling

Based on all the results, we propose the following mechanism for aster migration (see graphical abstract). The sperm aster grows in meiosis and moves off the cortex by pushing and potentially by cytoplasmic pulling. In interphase, cortical pulling brings the aster back to the cortex, despite the constant presence of cytoplasmic pulling. Finally, entry into mitosis triggers the migration of the spindle away from the cortex: cortical pulling stops at mitotic entry, MTs become more dynamic, and the mitotic apparatus moves by combined cytoplasmic pulling and cortical pushing. We designed a 2D agent-based stochastic computer model of aster migration based on the software *Cytosim* (Foethke et al., 2009) (Figure 7A, Movie 7). The parameters of model simulations were set according to past studies on different species (Sup Table S2), and adjusted to our data to match the timing of cell cycle and the cortical pulling activity. We verified the equivalence of the simulated and observed cytoplasmic pulling by comparing a simulation to latrunculin data (Figure S4 A and B, first row). Once an average reference simulation was established, we could remove specific forces to test the contribution of cortical pulling in the global scenario of aster migration (Figure 7), or to explore other situations (Figure S4). With a permanent cortical pulling the aster is quickly brought to the cell cortex in meiosis and did not seem to leave the cortex during the cell cycle (Figure 7A 7B, second row). Thus, in this condition the migration did not fit the control profile of aster migration as the aster was always further from the cell center than in control (Figure 7B, second row).

**Figure 7.**
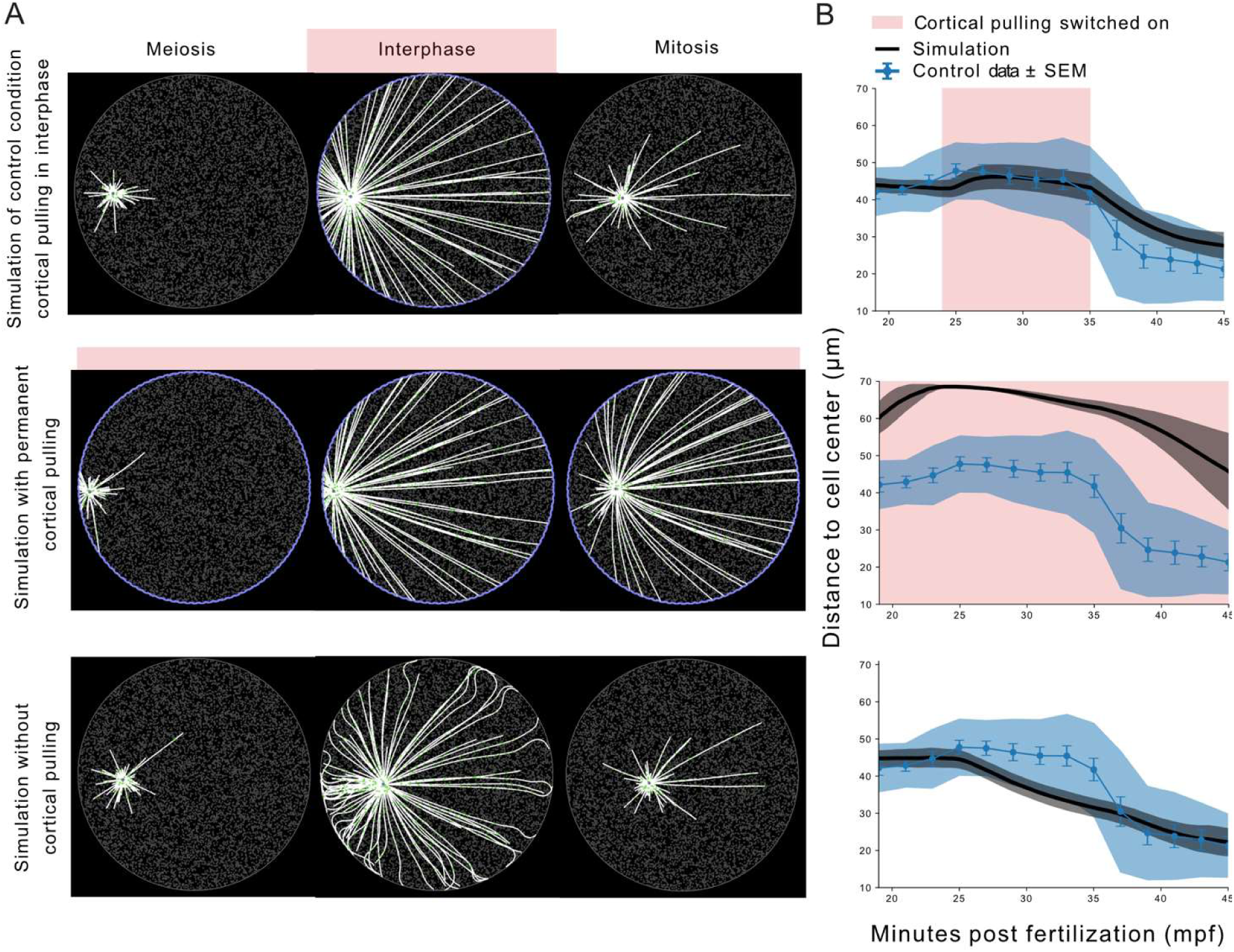
Cortical pulling dictates the pattern of aster migration. A. Stills from simulations testing the contribution of cortical pulling in the model of aster migration. The selected frames correspond to times of meiosis, interphase and mitosis in the simulation. The mitotic apparatus was simplified as an aster. The aster core (centrosome) is represented by a purple dot at the center of the MTs structure. MTs (in white) are set to become more stable in interphase. The purple cell border illustrates the activity of cortical dyneins, and thus of cortical pulling. The many gray dots represent fixed dyneins in the cytoplasm to reflect the presence of cytoplasmic pulling in the cell. They become green when attached to MTs. Three conditions were tested: a transient cortical pulling in interphase (first row) that serves as a reference simulation, a permanent cortical pulling (second row), and an absence of cortical pulling (third row). B. Comparison of the pattern of aster migration shown as the distance of the aster from the cell center through time, in control experimental data (blue curve), and in the simulations (black curve). The blue and gray shades represent +/-SD. The red margin represents the period of cortical pulling activity in the corresponding simulation. The graphics are aligned with A so that the simulation corresponding to the graphic appear on the same row.

In the complete absence of cortical pulling throughout the cell cycle (Figure 7, third row) the aster position was slightly more centered in interphase, and the overall aster centration hence reached the same distance from the cell center as in the average reference simulation (Figure 7 first and third rows). Indeed, in the absence of cortical pulling the aster centered as much as it did in the control dataset, however the centration began earlier, progressed at a constant velocity, and did not show the static phase prior to aster migration (Figure 7B, third row). The comparisons of simulations outputs to the data supported the scenario drawn from our results where aster centration is prevented by cortical pulling activity during interphase, and the inactivation of cortical pulling in mitosis permits aster movement. Only relying on simulations, we pushed the exploration of our model on questions still experimentally unanswered. These theoretical explorations suggested that cytoplasmic pulling is the main contributor to aster centration in mitosis compared to the pushing mechanism (Sup Figure S4A, S4B, bottom row). Finally, we used the simulation to examine a scenario where cortical pulling is nonuniformly inactivated at mitosis entry i.e., if cortical pulling is first turned off near the aster before being completely inactivated (Figure S4C). In this case, the aster centration during mitosis was faster than that obtained by simulating uniform inactivation (Figure 7 top) and the simulation best fitted the experimental data (Figure S4C).

## Discussion

A central question in the field of cell biology is how a cell divides into two equal sized daughter cells which relies on positioning of the mitotic apparatus to the cell center. The mechanism(s) involved in centering are still not fully resolved although they often depend on MT-based cortical pulling, cytoplasmic pulling or cortical pushing depending on the species, cell type and cell cycle stage. Centration has been studied extensively in *Xenopus,* sea urchin and *C. elegans* zygotes. In the large oocytes of *Xenopus* (circa 1mm) cytoplasmic pulling forces provide the force for aster centration. For example, by microinjecting a dominant negative fragment of the dynactin complex (p150-CC1) sperm aster centration was blocked (Wühr et al., 2010). Also, since MTs of the sperm aster are too short to reach the cortex on the opposite side of the zygote the primary mechanism for centration in *Xenopus* is cytoplasmic pulling in the direction of the longest MTs (Wühr et al., 2010). Sea urchins have smaller zygotes (circa 100µm) and even though MTs of the sperm aster are long enough to reach the opposite cortex, cortical pulling has not been reported to be involved and instead the two mechanisms reported to power sperm aster centration in sea urchin are based on cytoplasmic pulling (Tanimoto et al., 2016) and cortical pushing (Meaders et al., 2020). In *C. elegans* centration occurs during prophase in a dynein-dependent manner (Gönczy et al., 1999). In order to distinguish between cortical and cytoplasmic dynein, factors that recruit dynein to the cortex have been depleted. RNAi knockdown of goa-1/gpa-16 to deplete cortical dynein led to a higher velocity of sperm aster centration suggesting that cytoplasmic dynein was the primary force generating mechanism for centration (De Simone et al., 2018). In addition, since depletion of cortical dynein increased the velocity of centration cortical pulling forces were suggested to counteract the cytoplasmic forces that power centration of the sperm aster (De Simone et al., 2018).

The examples of sperm aster centration in *Xenopus* and sea urchin occur during interphase while in *C elegans*, primates (Asch et al., 1995; Hewitson and Schatten, 2002; Simerly et al., 2019) and ascidian (here) centration occurs at mitotic entry. Moreover, the position and geometry of the ascidian sperm aster, with long MTs extending into the zygote interior and short MTs directed towards the proximal vegetal cortex, represents a configuration that should favor cytoplasmic pulling to displace the sperm aster towards the zygote center. We were therefore curious about what prevented sperm aster centration until mitotic entry. A cell cycle link that triggers an increase in cortical pulling had been reported previously in *C. elegans* one cell stage embryos (Bouvrais et al., 2021; McCarthy Campbell et al., 2009; Redemann et al., 2010). An increase in posterior cortical pulling forces displaces the mitotic apparatus towards the posterior cortex during anaphase (Grill et al., 2003; McCarthy Campbell et al., 2009; Redemann et al., 2010). Indeed, the number of short-lived MT plus ends engaged in cortical pulling increased at the posterior pole of *C. elegans* one cell stage embryos at anaphase onset (Bouvrais et al., 2021). This increase in cortical pulling has been linked with the fall in cyclin-dependent kinase (CDK) 1 activity: either reducing the function of the proteasome, the APC (anaphase-promoting complex), or Cdc20 all delayed spindle displacement while inactivating CDK1 in prometaphase caused premature spindle displacement (McCarthy Campbell et al., 2009). Although these findings indicate that cortical pulling increases at anaphase (Keshri et al., 2020; Kotak et al., 2013) one key additional point may be that cortical pulling is less prominent when CDK1 activity is elevated. Here in the ascidian, we have noted a similar phenomenon whereby cortical pulling is elevated during interphase and reduced at mitotic entry when CDK1 activity increases.

In the ascidian *P. mammillata*, the sperm aster forms in the vegetal hemisphere of the zygote during meiosis II (Roegiers et al., 1995). Here we demonstrate that during meiosis II the sperm aster grows and remains roughly spherical as it moves slowly away from the proximal vegetal cortex (Figure 1). At this stage the sperm aster remains relatively small (approx. 1/3 zygote diameter) and located in the zygote vegetal hemisphere. Such a vegetal location and relatively small size ensures that sperm astral MTs do not reach the animal pole and thus interfere with the segregation of the meiotic chromosomes, which in smaller *C.elegans* zygotes is accomplished by preventing sperm aster growth during meiosis II (McNally et al., 2012). At entry into interphase astral MTs extend throughout the zygote and capture the female PN located at the animal pole (Figure 1). The female PN then migrates towards the center of the sperm aster during a short interphase (about 10 min) while the cortically-located and highly asymmetric sperm aster remains in position near the vegetal pole (Figure 1). Just prior to NEB, during prophase, the sperm aster begins abruptly to migrate accompanied by NEB and formation of a bipolar mitotic apparatus (Figure 1).

What triggers the switch to induce migration at entry into mitosis? First, we delayed entry into mitosis to determine if a causal relationship existed between entry into mitosis and sperm aster migration. Delaying entry into mitosis with the CDK inhibitor p21 prevented sperm aster migration (Figure 2). Next, we teased apart the relative contributions of cortical pushing, cytoplasmic pulling, and cortical pulling to determine which of these three mechanisms displayed a cell cycle-dependent change at mitotic entry that could explain how mitotic entry triggered sperm aster migration. Cortical pushing is present in Meiosis II, and is responsible for the initial aster displacement from the cortex (Figure 6). Even though it seems this mechanism is also active in mitosis (Figure 6B), its contribution to spindle migration at mitosis entry seems minor (Figure S4). We therefore focused on cytoplasmic pulling and cortical pulling. Cytoplasmic pulling can be visualized through the movement of three different endomembrane structures towards the center of the sperm aster: the female PN, endoplasmic reticulum and vesicles (Figure 3). Taking advantage of the opportunity to monitor a large number of vesicles (labelled with Cell Mask Orange) we quantified the movement of the cytoplasmic vesicles to determine whether there was a measurable difference in cytoplasmic pulling between interphase and mitotic entry (Figure 3 and Figure S2). The data demonstrated that vesicles transport was unchanged between interphase and mitotic entry, suggesting a constant cytoplasmic pulling. We then sought to determine whether cortical pulling was more prominent during interphase or mitosis. To do so we exploited the membrane invagination assay following weakening of the cortex (Godard et al., 2021; Redemann et al., 2010; Rodriguez-Garcia et al., 2018). Interestingly, we noted that cortical pulling was greater during interphase than mitotic metaphase (Figure 4). We therefore devised a 2-cell stage assay to determine whether cortical pulling was a feature of interphase and switched off at mitotic entry. By either delaying mitotic entry with p21 or blocking exit from metaphase with Δ90 cyclin B in one sister cell (Figure 4) we demonstrated that cortical pulling is elevated during interphase and switched off at mitotic entry when CDK1 activity is elevated. This observation could explain the switch-like behavior in migration and suggests that it is the increase in CDK1 activity at mitotic entry that switches off cortical pulling thus liberating the sperm aster from its cortical tethers facilitating centration. Moreover, these data develop further the findings from *C. elegans* where cortical pulling was decreased when CDK activity was elevated (McCarthy Campbell et al., 2009).

By using the software *Cytosim* we tested whether cortical pulling could prevent aster migration mediated by cytoplasmic pulling. Simulations showed that cortical pulling was capable of preventing migration caused by long MT-mediated cytoplasmic pulling and re-enforce the data showing that the sperm aster does not migrate during interphase when cortical pulling is elevated (Figure 7). This supports the idea that switching off cortical pulling at mitotic entry is necessary for sperm aster migration. Furthermore, the simulation indicated that a total absence of cytoplasmic pulling prevented aster migration in mitosis (Figure S4). Overall, these findings demonstrate that mitotic apparatus migration in the ascidian occurs at mitotic entry which causes the switching off of cortical pulling while cytoplasmic pulling and pushing remain active. It would be interesting to examine the relationship between mitotic entry and sperm aster migration in primate zygotes to determine whether cortical pulling is also switched off at mitotic entry.

## Acknowledgments

We thank members of the Turlier and McDougall groups for technical advice and discussion. We are grateful to the Imaging Platform (PIM) and animal facility (CRB) of Institut de la Mer de Villefranche (IMEV), which is supported by EMBRC-France, whose French state funds are managed by the ANR within the Investments of the Future program under reference ANR-10-INBS-0, for continuous support. This work was supported by a collaborative grant from the French Government funding agency Agence National de la Recherche to McDougall (ANR ‘‘MorCell’’: ANR-17-CE 13-0028), by a MITI award from the CNRS (Modélisation du vivant) by an Assemble + grant 9632 to Burgess, and by Sorbonne Université which provided a doctoral stipend. HT has received funding from EMBRC-France (AAP Découverte 2020) and from the European Research Council (ERC) under the European Union’s Horizon 2020 research and innovation program (Grant agreement No. 949267).

## Supplementary Figures and Table

**Supplementary Figure S1.**
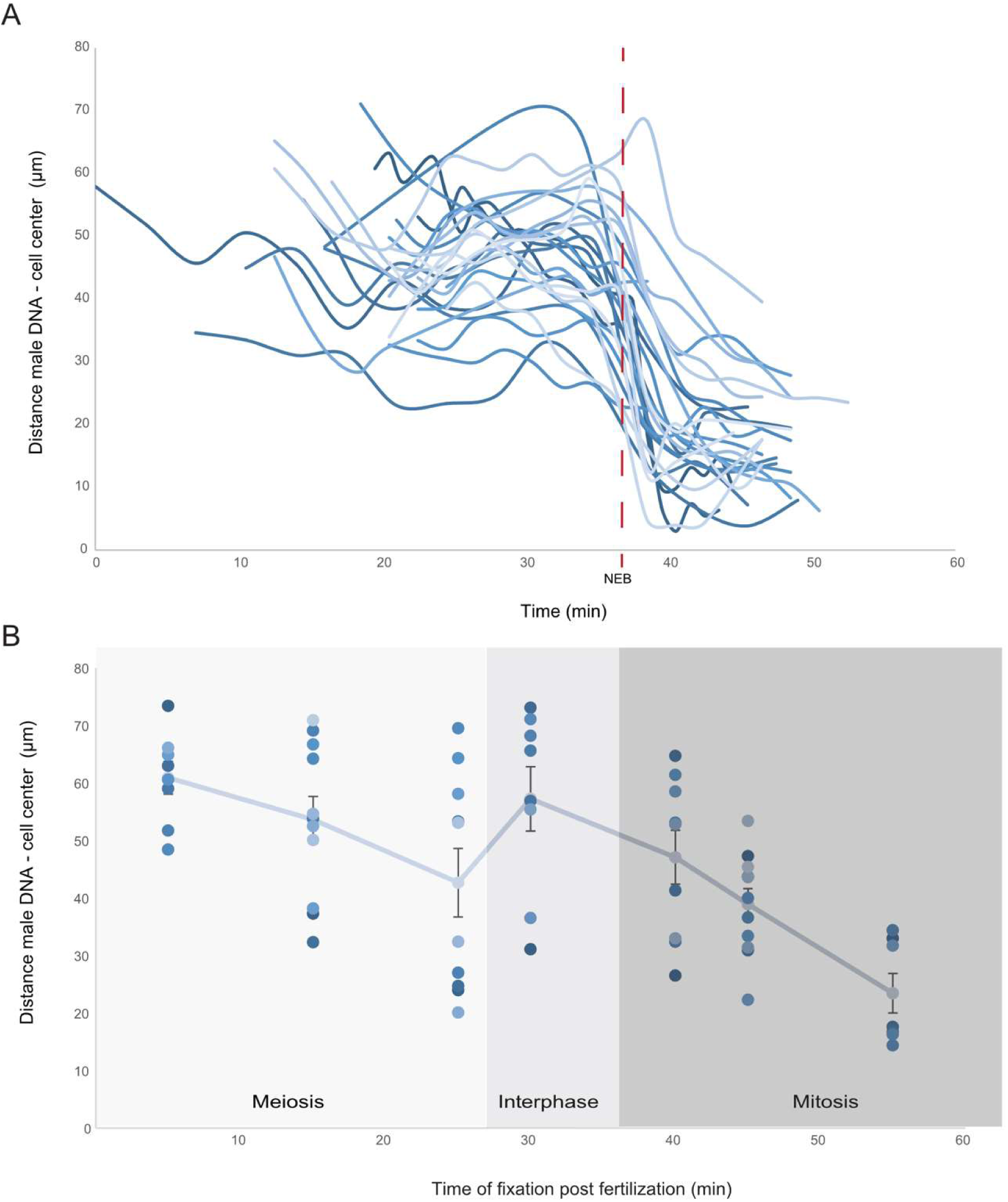
Live and fixed data displaying sperm aster position following fertilization. A. Sperm aster position relative to the cell center as a function of time in live zygotes. All individual zygotes exploited in Figure 1 are displayed and aligned with respect to their time of NEB, indicated by a red dotted line. B. Sperm aster position relative to the cell center in fixed zygotes.

**Supplementary Figure S2.**
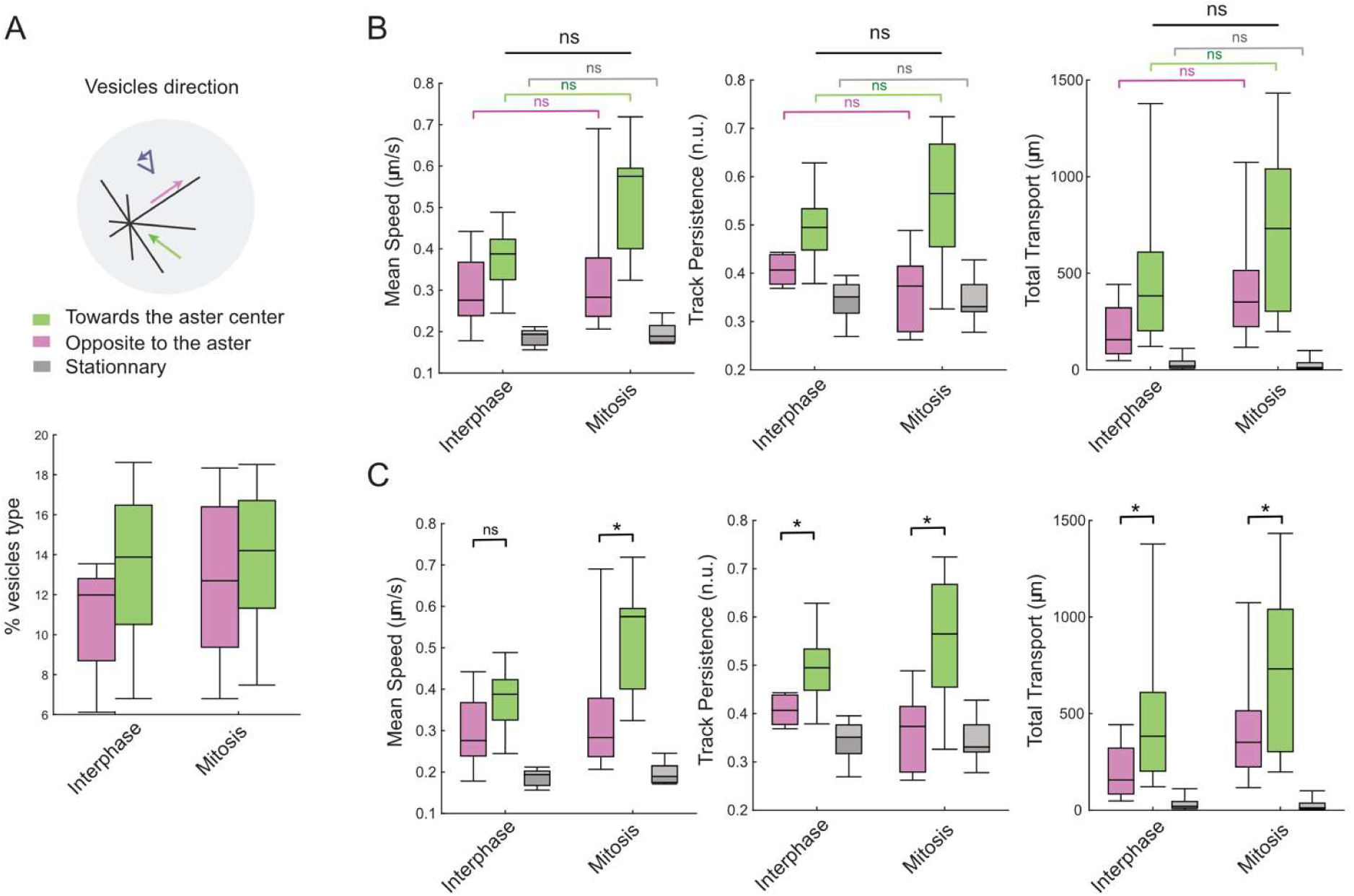
Vesicle movement parameters. A. Schematic showing vesicles either moving towards (green), away from (pink) the sperm aster center, or stationary (gray arrow). The graph shows the mean percentage of the 2 types of moving vesicles in interphase and in mitosis. The majority of vesicles are stationary (not shown in the graph). B. Statistical comparison of interphase versus mitosis for vesicle movement parameters: mean speed, track persistence, and total transport. Track persistence is the distance traveled towards the aster over the total trajectory of the vesicle. Total transport is the cumulative distance travelled by all vesicles during the 3 minutes time period. No significant difference was found. n=8 embryos, mean number of vesicles tracked per embryo = 1129 in interphase, 1734 in mitosis. Paired Wilcoxon test. C. Statistical comparison of vesicles moving towards or away from the aster center. From the same dataset as presented in B. Paired Wilcoxon test.

**Supplementary Figure S3.**
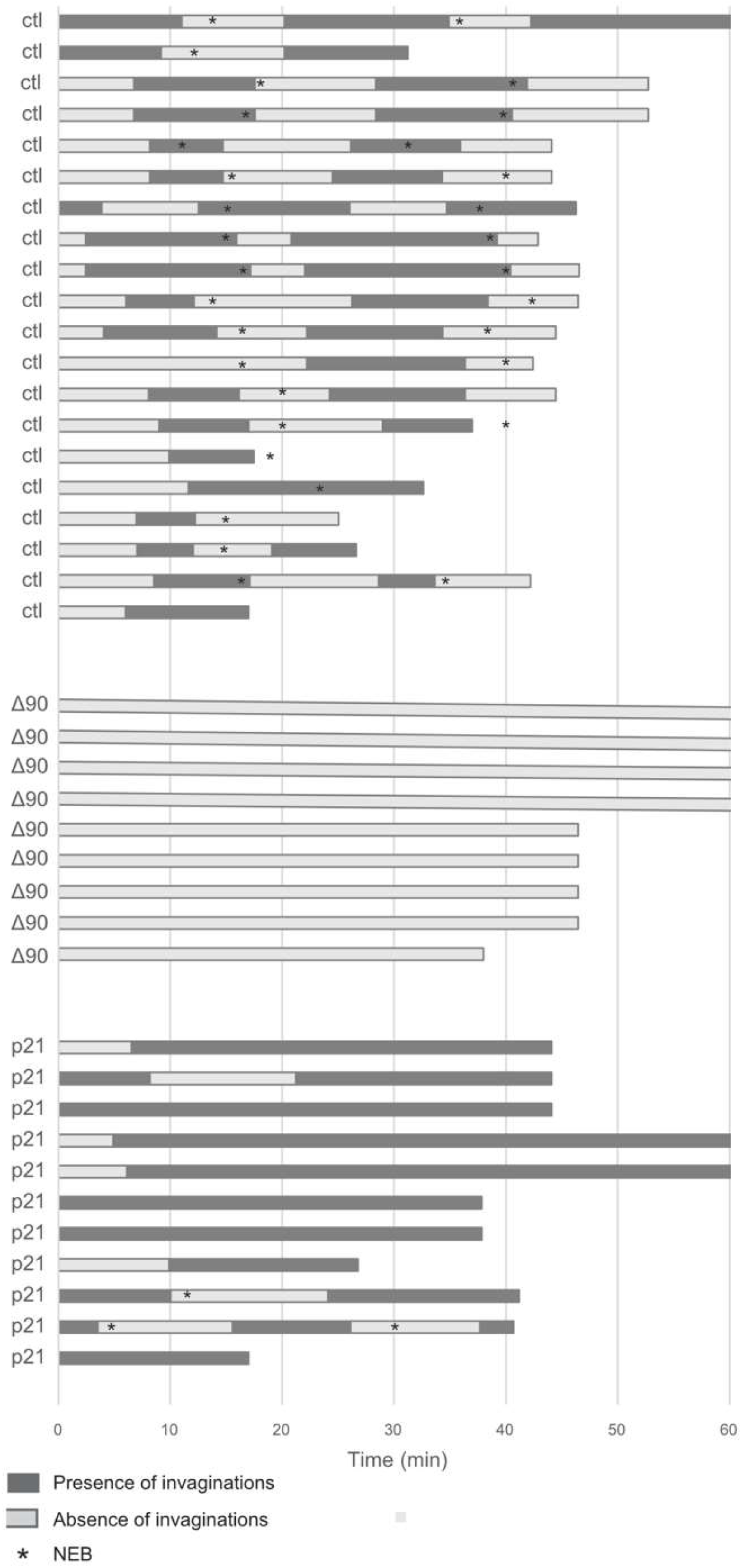
Effect of cell cycle inhibitors p21 and Δ90 cyclin B on the presence of membrane invaginations. Compiled data showing phases of membrane invaginations as a function of time. 2-cell stage embryos in which membrane was fluorescently labelled (by previous injection of PH::tomato mRNA or incubation in CellMask Orange) were injected with cell cycle inhibitors then treated with cytochalasin and invaginations were imaged by confocal microscopy. Each line represents 1 cell. Dark bars represent the presence of invaginations and light bars the absence of invaginations for control cells (n=20), cells injected with Δ90 Cyclin B protein which blocks in metaphase (n=9) or cells injected with p21 protein which prolongs interphase (n=11). * represents time of NEB. Also see Movies 5A and 5B.

**Supplementary Figure S4.**
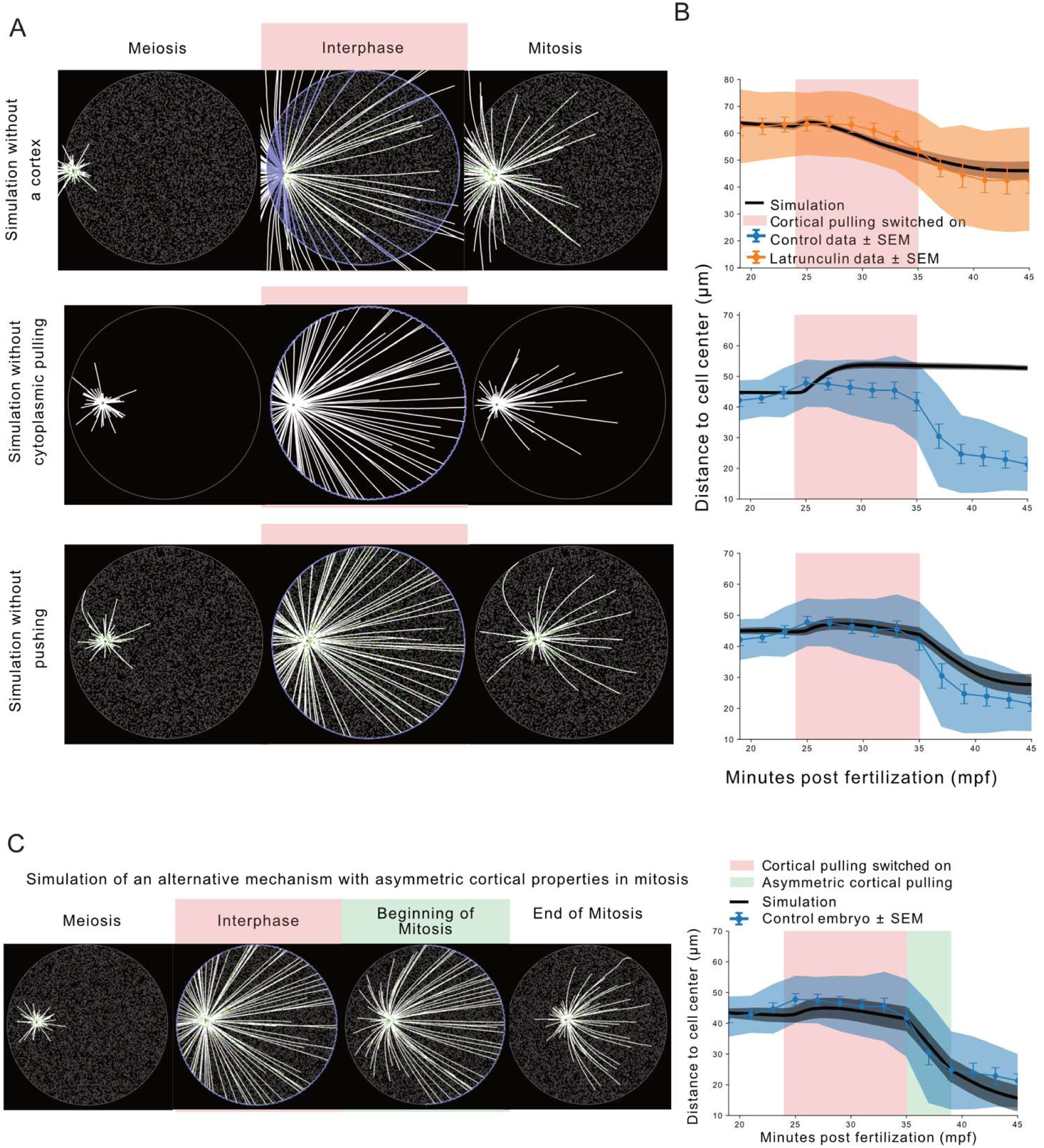
Contribution of pushing and cytoplasmic pulling in aster centration. A. Stills from simulations testing the contribution of pushing and cytoplasmic pulling in the aster migration pattern. Like previously (Figure 7), the selected frames correspond to times of meiosis, interphase and mitosis in the simulation. The aster core (centrosome) is represented by a purple dot at the center of the MTs structure. MTs are in white, they are set to become more stable in interphase. The purple cell border illustrates the activity of cortical dyneins, and thus an active cortical pulling. The many gray dots represent fixed dyneins and thus the presence of cytoplasmic pulling in the cell. They become green when attached to MTs. Three conditions were tested. To test the contribution of cytoplasmic pulling alone (first row), as performed experimentally by destroying the cell cortex (Figure 5), the pushing was removed by allowing MTs to ignore boundaries, and the cortical pulling was drastically reduced. Because MTs ignore boundaries, their plus end is often located outside the cell at interphase onset, so cortical dyneins were allowed to bind on all MTs, not only their plus end. The cortical motors were still allowed to bind MT but, as seen with the phenomenon of membrane invagination (Figure 4), they were pulled towards the MT rather than strongly attached to the cell boundary. The second row shows the inhibition of the cytoplasmic pulling, where the fixed cytoplasmic dyneins were removed. Finally, in the third row we tested the inhibition of the pushing mechanism without affecting the cortical pulling, which was experimentally impossible with the tool used in this study. MTs were not allowed to ignore the cell boundary, thus the inhibition of pushing was done by preventing the MT to have a grip on the cell boundary. B. Aster migration, as the distance of the aster from the cell center through time, in simulations (black curve) compared to control experimental data (blue curve) or latrunculin-treatment (orange curves). The blue, orange and grey shades represent +/- SD. The red rectangle shading indicates the period of cortical pulling activity in the corresponding simulation. Each graph in B is aligned with the corresponding simulation in A. C. Simulation exploring a mechanism of aster migration where, at mitosis entry, the cortical pulling is first turned off near the aster before being completely inactivated. The resulting migration (black curve) is compared to control data (blue). The green margin shows the moment of asymmetric cortical pulling.

**Supplementary Table S1.** Injection of p21 prolongs interphase. Analysis of the effect of p21 on the duration of cell cycle phases. Comparison between control and p21-injected zygotes for the duration between pronucleus formation to NEB and between NEB and cytokinesis. p21 significantly prolonged first interphase since the average duration from PN formation to NEB went from 13 min to 28 min. n is displayed together with the SD.

| Cell Cycle events | Control |  |  | p21-injected |  |  |
| --- | --- | --- | --- | --- | --- | --- |
|  | min post fertilization | min after PN formation | n | min post fertilization | min after PN formation | n |
| PN formation | 25 ± 2.8 | 0 | 22 | 34.9 ± 11.3 | 0 | 17 |
| NEB | 37.5 ± 4.2 | 12 | 23 | 65.8 ± 14.2 | 30 | 16 |
| Cytokinesis | 48 ± 4.5 | 23 | 24 | 65 ± 5.1 | 29 | 9 |
|  | min |  |  | min |  |  |
| Time between PN formation & NEB | 12.7 ± 2.6 |  | 20 | 26.9 ± 2 |  | 15 |
| Time between NEB & Cytokinesis | 10 ± 2 |  | 23 | 11.3 ± 1.8 |  | 9 |

**Supplementary Table S2.**
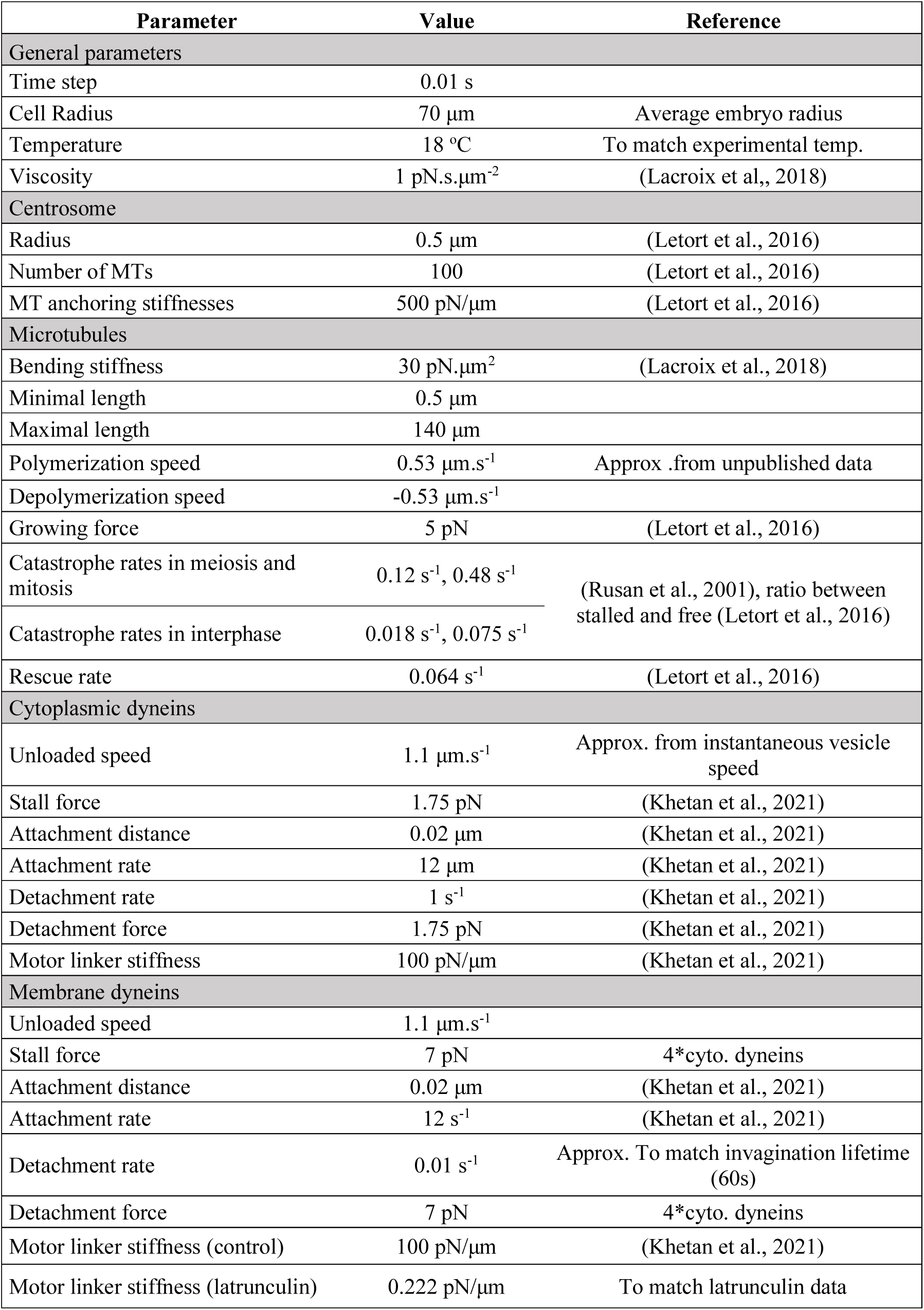
Parameters used in the simulations related to Figure 7 and Figure S4. The *Cytosim* parameters used to create the simulations are listed in the table along with the corresponding references.

## Supplementary Movies

**Movie 1.** Confocal timelapse of aster formation and migration.

Confocal z-sections from a xyzt series showing microtubules in a live zygotes injected with Ens::3GFP. Scale bar is 50µm. http://movincell.org/medias/609

**Movie 2.** Female PN migration.

Confocal z-section from a timelapse series showing microtubule plus ends and associated brightfield showing female pronuclear migration. Microtubules are labelled with EB3::GFP. http://movincell.org/medias/610

**Movie 3.** Cell Mask Orange labelled vesicles accumulating at the sperm aster

A. Confocal timeseries with images collected every second in one z plane. The sperm aster is at the bottom and accumulates red fluorescence as vesicles become localized to the sperm aster. http://movincell.org/medias/611

B. Confocal timeseries with images collected every second in one z plane. Here the zygote was compressed. The sperm aster is at the bottom and accumulates red fluorescence as vesicles become localized to the sperm aster. The female pronucleus in compressed zygotes formed karyomeres which can be seen migrating towards the sperm aster. http://movincell.org/medias/612

**Movie 4.** Cycling invaginations in the zygote

Zygote treated with latrunculin and labelled with Cell Mask Orange. No membrane invaginations are present during meiosis II. Membrane invaginations first become visible during interphase. The membrane invaginations are subsequently lost at mitotic entry. Scale bar = 50µm. http://movincell.org/medias/613

**Movie 5.** Membrane invagination during interphase in 2 cell stage embryo with one cell injected with either p21 or Δ90 Cyclin B.

A. Confocal timeseries of 2-cell embryo treated with latrunculin and labelled with Cell Mask Orange (rendered cyan). Membrane invaginations are present in the non-injected cell during interphase and are permanently present in the p21::GFP-injected cell (rendered magenta, right cell) during the prolonged interphase. Scale bar is 50µm. http://movincell.org/medias/614

B. Confocal timeseries of 2-cell embryo treated with latrunculin and labelled with Cell Mask Orange (rendered cyan). Membrane invaginations are present in the non-injected cell only during interphase, and not in Δ90 Cyclin B::GFP-injected cell (rendered magenta, cell on the right). Scale bar is 50µm. http://movincell.org/medias/615

**Movie 6.** MT pushing on plasma membrane in meiosis/interphase/mitosis.

Confocal timeseries of the outward membrane protrusions in the zygote. Microtubules are labelled with Ensconsin::3GFP, the plasma membrane is labelled with Cell Mask Orange and the zygote is treated with latrunculin. http://movincell.org/medias/616

**Movie 7.** Simulation of the aster migration

2D simulation of the aster migration from mid-meiosis to mitosis. The aster core (centrosome) is represented by a purple dot at the center of the MTs structure. MTs (in white), are set to become more stable in interphase and when they touch the cortex. The purple cell border illustrates the activity of cortical dyneins, and thus an active cortical pulling, also indicating interphase. The many grey dots represent fixed dyneins and thus the presence of cytoplasmic pulling in the cell. They become green when attached to MTs. http://movincell.org/medias/617

## Materials and Methods

### Biological material

*Phallusia mammillata* adult animals were collected in Roscoff or Sète and kept at 16°C in the aquaria of the “Centre de Ressources Biologiques” (CRB) of the Institut de la Mer à Villefranche (IMEV) which is an EMBRC-France certified service (see https://www.embrc-france.fr/fr/nos-services/fourniture-de-ressources-biologiques/organismes-modeles/ascidie-phallusia-mammillata). The gametes were collected by puncturing separately the oviduct and the sperm duct. The sperm was kept dry at 4°C and could be used for fertilization up to 1 week after collection. Oocytes were used the day of collection after undergoing dechorionation by incubation in 0.1-0.2% trypsin in micro-filtered natural sea water (MSFW) at 19°C for 90 minutes, and subsequent washes in MSFW supplemented with 5mM TAPS (tris(hydroxymethyl)methylamino] propanesulfonic acid) pH 8.2. All the subsequent manipulations of live embryos were performed in MSFW 5mM TAPS, using pipette tips, dishes, slides and coverslips coated with 0.1% gelatin and 0.1% formaldehyde (Sardet et al., 2011). For fertilization, a small volume of activated sperm (circa 5µl) was added to the oocytes in a 5ml petri dish. To activate the sperm, 6 µL of dry sperm was incubated for 20 minutes at 19°C in 500µL of MSFW pH9.2. Time post fertilization was measured starting when the oocyte first showed a shape change. For fixation of fertilized cultures, the time of fertilization was determined when about 30% of the oocytes showed the first deformation.

### mRNA synthesis and injections

Synthetic mRNAs for microinjection were prepared using the mMESSAGE mMACHINE T3 kit (Ambion), from plasmids containing the gene of interest (EB3::3GFP, Ensconsin::GFP, PH::GFP, Histone H2b::Rfp1, Venus::Reticulon) flanked by a T3 promoter and a polyA tail. mRNA yield was estimated by spectrophotometry. The mRNAs were stored at high concentration (>10µg/µL) in 1 μl aliquots at −80 °C, then thawed and diluted in distilled water for use. The mRNAs were micro injected into dechorionated oocytes transferred to small glass wedges mounted onto 400 µl Perspex mounting chambers designed for horizontal microinjection (see detailed protocols in McDougall et al., 2014). mRNAs were injected at a pipette concentration of 5-6 μg/μl (injection volume is ∼1–2% volume of the egg) using a high pressure system (Narishige IM300). mRNA-injected oocytes were left for 5 h or overnight before fertilization and subsequent confocal imaging.

### Protein purification and injections

The p21::mCherry protein (p21 is the cyclin dependent kinase inhibitor 1a) was injected to arrest the cell cycle in interphase. The human p21 protein fused with mCherry was cloned in a pET11a vector with 6 His-tag, and purified with a silica-based resin column (MACHEREY-NAGEL, protino Ni-IDA). Aliquots were flash frozen in liquid nitrogen and stored at –80°C. The protein was injected at a concentration of 30mg/ml.

The construction and synthesis of the human Δ90 cyclin B1::GFP plasmid has been described previously (Levasseur and McDougall, 2000). The Δ90cyclin B::GFP fusion protein was stored at –80°C and was injected at a final concentration of approximately 11 mg/ml.

The microinjection system described above for mRNAs injection was also used for protein injection. Protein-injected oocytes were left for 45min at 18°C before fertilization and subsequent confocal imaging.

### Confocal microscopy imaging

All imaging experiments were performed at 19°C using a Leica TCS SP8 inverted microscope fitted with Hybrid detectors and 40×/1.1NA water objective lens. To image aster migration, each fertilized egg or 2-cell stage embryo was scanned by 4D live imaging of a whole embryo (xyzt) with a frame rate of at least 1 image every 210s. The imaging parameters were adapted to each fertilized egg: z step was between 0.5 and 2µm and time step between 1 min and 3.5 min. For fixed samples, z-stacks were acquired at step size of 0.5 µm.

### Invagination experiments

Eggs were fertilized in MSFW and transferred after observing the first deformation into a solution of MSFW TAPS containing 5µM of latrunculin B (Sigma Aldrich) diluted from a 10mM stock solution (in DMSO). The embryos were then mounted on a slide in the latrunculin SW solution. The zygote plasma membrane was visualized either by microinjection of the PH::Tomato mRNA, or by addition of the membrane dye Cell Mask orange (Thermofisher, Invitrogen) at 1:1000. For controls, zygotes labeled with PH::Tomato or Cell Mask were treated with DMSO at a dilution of 1:1000.

In the case of the embryo injected at the two-cell stage, the fertilized eggs were left to develop at 16°C in MFSW until they started cleaving. Then they were mounted in the injection chamber, and injected with p21 or Δ90cyclinB proteins as soon as the division finished. When two or three embryos were injected they were immediately transferred in MFSW with 5µM of Cytochalasin B (Sigma Aldrich) diluted from a 10mM stock solution in DMSO, and mounted on a slide in this solution for imaging.

### Quantification of the number of invaginations

To image membrane invaginations in zygotes or 2-cell stage embryos, 25 µm-thick stacks of confocal images (dz=2.5 µm) were acquired every 10 seconds in Cell Mask Orange stained zygote at around 2-4 minutes after latrunculin incubation. The number membrane invaginations was counted manually by counting invaginations present at 2-5 µm from the plasma membrane.

Membrane invaginations were imaged in two-cell embryos 4-5 minutes after incubation in cytochalasin B at a frequency of 1 z-stack every 30 seconds to 1 stack every 2 minutes. The presence or absence of invagination was scored manually on a z-projection image and cell cycle state of the cells were defined as the 15 minutes following NEB.

### Quantification of aster migration

The quantification of the distance between the center of the zygote and the DNA throughout the cell cycle was performed in three steps, using Fiji (Schindelin et al., 2012).

First, the center of mass of zygotes were determined at each time point. To do so, a Gaussian Blur (sigma =2) was applied to each xyz stack of the timelapse movie. Stacks were then made binary with the “Triangle” method of the “Auto Threshold” function. The Fiji plugin “3D Roi Manager” (Ollion et al., 2015) created objects from the binary stacks, and output their center of mass. This method was verified by comparing the center of mass of the 3D zygote to the centroid of the 2D equatorial section with widest diameter.

Secondly, using the Fiji “Point tool” and the “measure” function, the xyz coordinates of the DNA label were obtained at each time point. When DNA was not labeled (in p21-injected embryos), or weak (at NEB), DNA position was approximated to be at equal distance between the two spindle poles, or at the center of the aster when the spindle was not yet formed. The DNA label was chosen over the MT label to measure the aster migration because centrosome duplication occurs before the centration of the spindle, therefore while the spindle as a whole centers, each spindle pole starts centering and then diverges from the cell center to center the DNA.

Finally, the distance between the DNA coordinates and the center of mass was computed using the 3D Pythagorean theorem: d=√((x2−x1)^²^+(y2−y1)^²^+(z2−z1)^²^).

### Quantification of the vesicle traffic

To image vesicle trafficking, eggs were fertilized, washed once, and immediately transferred to a GF-coated slide/coverslip in a MFSW containing Cell Mask orange (1/1000). Then, 2D images acquired every second for 3 min. The imaged plane was selected to contain the center of one aster.

To measure relative movements of the vesicles, we combined 3 approaches. 1) For vesicles, we used the Fiji tool TrackMate with LoG particle detection and simple LAP tracker (Tinevez et al., 2017). 2) For aster localization we wrote (using Matlab) a manual periodic tracking, with interpolation for intermediate time frames. 3) For cell contour, we developed another Matlab algorithm based on threshold optimization to extract the cell contour. We combined information from those tools to quantify movement.

For each vesicle track, we measured the relative path with respect to the aster. In more detail, we defined a radius from the aster center to the centroid of the track, which naturally crosses the cell contour. On this radius, we projected the path to estimate the radial component of the vesicle movement. We also measured the temporal evolution of the contour. To take into account the cell deformation and its impact on vesicle movement, we subtracted from each relative path a yield drift. Considering an elastic behavior on the aster-contour axes, the yield drift of a vesicle at a radius *r* was defined as follows:

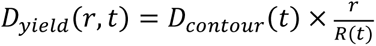

with *R(t)* the distance of the cell contour from the aster center. To segregate vesicles just endocytosed, stagnant below the membrane, from the vesicle moving on the aster, we kept only the tracks 10µm away from the cell contour. Based on the path projection on the radius and its orientation (positive if the last position is further from the aster than the first position), we then sorted the tracks as going toward (retrograde) or away from (anterograde) the aster. Vesicles with path projection smaller than 1µm were defined as stationary (note that this category includes the static vesicles and vesicles moving orthogonally to the radius).

### Modeling

Agent-based simulations were performed in 2D using a custom version of the software *Cytosim* - www.cytosim.org (Foethke et al., 2009) with the parameters provided in Table S2. *Cytosim* is a stochastic simulation engine that includes constituting elements of the cytoskeleton. It has already been used to study spindle and centrosome dynamics and position (Khetan and Athale, 2016; Lacroix et al., 2018; Letort et al., 2016).

#### Aster and microtubules

MTs are modeled as worm-like chains, characterized by a bending stiffness (see Table S2) and inextensibility. MTs can (de)polymerize and their instability is modeled by a stochastic alternation of growing and shrinking phases. The plus-end polymerizes until a catastrophe happens, starting a shrinking period. The aster is modeled as a bead, from which MTs are nucleated. MT minus-ends are anchored to the aster while their plus-end are directed outward from the aster.

A pushing force is generated by MTs’ plus end polymerizing against the edge of the cell. When MTs push strongly on the cortex, polymerization is slowed down and they have a higher chance to undergo a catastrophe. Previous work (Letort et al., 2016) suggests that pushing cannot center the aster if MTs can glide along the cell membrane. In their work, gliding was prevented by pinning MTs’ tips to the point where they first reached the edge of the cell. Pinning and cortical pulling were not activated at the same time though, as pinning would make cortical pulling inefficient. To prevent MT gliding along the cell cortex, we didn’t model control cells as a proper circle, but as a crenelated polygon, representing the actomyosin cortex. Once a tip enters an alcove, it cannot slide anymore, as if it were stuck by intertwined actin filaments.

#### Dynein distributed in the cytoplasm

Dyneins are placed at random positions in the cytoplasm. When a MT comes close, a dynein can bind to the MT and starts moving towards its minus end. As in previous work cited above, a spring-like force pulls the dyneins back to their assigned position when they are displaced. The dynein’s velocity depends on the load and the projection of the restoring force along the direction of the MT.

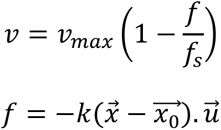

*v_max_* is the unloaded speed, ie the speed when there is no applied force. *f_s_* is the stall force, the maximal force the dynein can withstand before it stops moving. *k* is the spring stiffness, 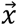 the position of the dynein, 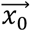 the rest position it has been assigned and 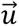 the direction of the MT.

#### Control of MT dynamics by cortical dynein to generate forces

If cortical pulling is implemented by usual dynein, it often makes the aster spin around the cell. This is due to the fact that MTs tend to align with the cell’s edge, as more and more cortical dynein becomes attached to the MTs. The pulling force becomes higher and higher and the aster spins around. *In vitro* experiments suggest that dynein placed in front of a rigid barrier can control MTs’ dynamics (Laan et al., 2012). Such dyneins trigger a catastrophe when they bind to the end of a MT, and regulate its shrinking speed thereby generating pulling forces. We modified *Cytosim* to implement such a behavior: cortical dynein works like classical molecular motors except that they can only bind to a MT end, and they chew the MT as they move forward. Like cytoplasmic dyneins, they have a stall force and the load comes from a spring linking the dynein to its original position. However, unlike cytoplasmic dyneins, cortical dyneins cannot move backwards as it would imply the MTs polymerized again.

#### Code availability and simulation reproducibility

The custom version of *Cytosim* with this interaction implemented is available at https://gitlab.com/gslndlb/cytosim, in the branch *dynein_chew*. All configuration files used are in the cym folder.

### Statistics and diagrams

Statistical tests and graphics were performed using the libraries rstatix, tidyverse, dplyr, ggpubr, and ggplot2 from R software (R Studio, 2020) as well as Microsoft Excel (2013). Tests are provided in the figure legends. Diagrams were created with BioRender.com

### Data availability

All main figures are supplied with data used to generate the figures.

## Notes

### Competing Interest Statement

The authors have declared no competing interest.

